# Cytotoxic CD4^+^ T cells in the bone marrow compromise healthy ageing by enhancing granulopoiesis

**DOI:** 10.1101/2024.01.26.577360

**Authors:** Enrique Gabandé-Rodríguez, Gonzalo Soto-Heredero, Elisa Carrasco, Carlos Anerillas, José Ignacio Escrig-Larena, Sandra Delgado-Pulido, Isaac Francos-Quijorna, Manuel M. Gómez de las Heras, Álvaro Fernández-Almeida, Eva María Blanco, Elia Winand-Osete, Virginia Zorita, Jorge Martínez-Cano, Amanda Garrido, Rafael de Cabo, Santos Mañes, Myriam Gorospe, María Mittelbrunn

## Abstract

Neutrophils are the most abundant leukocytes in the blood, with numbers further increasing with age. Despite their essential role as a primary line of defense, neutrophils can contribute to tissue damage and age-related diseases ^1^ and a high neutrophil-to-lymphocyte ratio predicts all causes of mortality in the elderly ^2–5^. However, the precise mechanisms driving enhanced neutrophil generation during ageing remain poorly understood. Here, we show that a subset of CD4^+^ T cells with a cytotoxic phenotype (CD4^+^ CTLs) producing the chemokine CCL5 and harbouring dysfunctional mitochondria, infiltrate the bone marrow and induce granulopoiesis in aged mice. During ageing, hematopoietic stem cells upregulate CCR5, the primary receptor for CCL5, and its deficiency limits the T cell-mediated induction of granulopoiesis and neutrophil output. Treatment with the FDA-approved CCR5 inhibitor Maraviroc decreases granulopoiesis and lowers the levels of circulatory and tissue-infiltrating neutrophils, ameliorating multiple ageing biomarkers and improving functional outcomes in aged mice. These findings suggest that age-associated alterations in T cells reduce health outcomes by remodelling the bone marrow niche and enhancing neutrophil generation. Consequently, interventions to disrupt the interplay between T cells and hematopoietic stem cells hold substantial therapeutic potential to ameliorate age-associated diseases.

## Main

During ageing, impaired immune cell function increases the risk of suffering cancer and infections. Additionally, faulty immune cells sustain inflammageing ^6^, a persistent chronic inflammation that exacerbates several age-associated pathologies including cardiovascular disease, diabetes, chronic kidney disease, non-alcoholic fatty liver disease, autoimmune and neurodegenerative disorders, jointly representing the leading causes of disability and mortality worldwide ^7,8^.

Despite their relatively short lifespan, neutrophils are the most abundant leukocytes in the blood. Upon encountering danger signals, neutrophils rapidly migrate to target tissues to efficiently eliminate pathogens employing various mechanisms such as phagocytosis, degranulation, or suicide to release DNA in neutrophil extracellular traps. However, increased numbers of hyper-responsive neutrophils may contribute to tissue damage during ageing ^9^. For example, neutrophils engaged in reverse transendothelial migration re-enter the circulation from inflamed aged tissues and disseminate to the lungs, causing vascular leakage and remote damage ^10^. Moreover, tissue-infiltrating neutrophils have the capacity to induce paracrine senescence by triggering telomere damage through the release of reactive oxygen species ^11^.

The number of circulating neutrophils increases with age, in part due to the skewing of haematopoiesis towards the production of myeloid precursors at the expense of lymphoid progenitors ^12^. This imbalanced differentiation ultimately results in an increased neutrophil-to-lymphocyte ratio (NLR) in the circulation. Remarkably, a high NLR serves as a strong predictor of mortality in virtually all age-associated diseases ^2–5^. Therefore, understanding the mechanisms driving enhanced granulopoiesis and a heightened NLR during ageing may critically change the way multiple age-associated diseases are managed.

Recent reports indicate that T cells infiltrate the bone marrow (BM) and eventually lead to increased granulopoiesis and circulatory neutrophils under acute psychological stress, prolonged starvation, or autoimmune disease^13–16^. Here, we report that ageing is associated with increased numbers of CD4^+^ CTLs in the BM. By paracrine signalling to adjacent aged hematopoietic stem cells (HSCs), CD4^+^ CTLs induce granulopoiesis, rising the levels of peripheral neutrophils and augmenting the NLR through a targetable CCL5/CCR5 axis, ultimately contributing to inflammageing and tissue senescence.

## Results

### Accumulation of CD4^+^ T cells in the bone marrow enhances granulopoiesis and increases CXCR4^hi^ CD62L^lo^ neutrophils in old mice

To understand how ageing influences the generation and fate of neutrophils, we performed multiparametric spectral flow cytometry to analyse the percentage of the different blood populations in old (24 months old) versus young (3 months old) mice. Uniform manifold approximation and projection (UMAP) representation showed increased numbers of circulating neutrophils and monocytes together with reduced circulating lymphocyte numbers (Fig. 1a and Extended Data Fig. 1a,b), resulting in an increased NLR (Fig. 1b). Of note, we observed a higher percentage of pro-inflammatory circulating CXCR4^hi^CD62L^lo^ neutrophils and a decreased percentage of CXCR4^lo^CD62L^hi^ naïve neutrophils ^17,18^ in old mice (Fig. 1c). Compared with CXCR4^lo^CD62L^hi^ neutrophils, CXCR4^hi^CD62L^lo^ neutrophils displayed a pro-inflammatory profile characterized by the upregulation of ICAM-1, TLR-4, CD11b, CD66a or CD101 and the downregulation of CXCR2 (Extended Data Fig. 1c). The increased numbers of circulating CXCR4^hi^CD62L^lo^ neutrophils correlated with an increased presence of neutrophils exhibiting a pro-inflammatory phenotype in the spleen (Extended Data Fig. 1d), and increased infiltration in the kidney, lung and liver in old mice compared with young mice (Extended Data Fig. 1e,f). To study whether these changes in the phenotype of neutrophils occurred by exposure to an aged environment, we performed heterochronic adoptive transfer experiments of peripheral blood leukocytes from young mice (CD45.1) to either young or old recipient mice (CD45.2). Four hours later, we analysed the phenotype of the transferred CD45.1 neutrophils in the blood (Fig. 1d). Transferred neutrophils acquired a CXCR4^hi^CD62L^lo^ profile more rapidly in old than in young recipients (Fig. 1e,f), supporting a cell-extrinsic induction of this phenotype.

**Figure 1.**
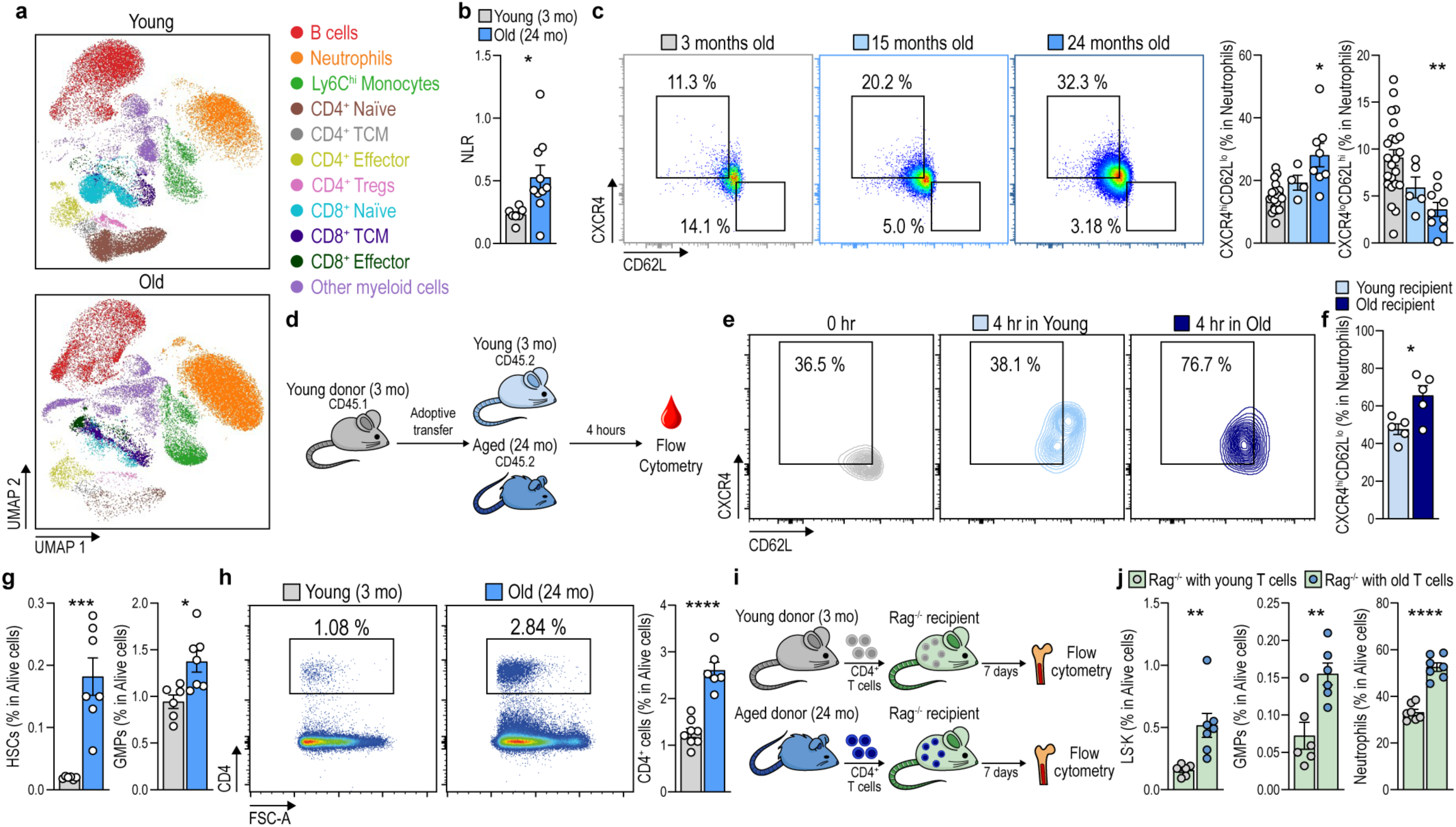
Accumulation of CD4^+^ T cells in the bone marrow leads to enhanced granulopoesis and increased levels of pro-inflammatory neutrophils in aged mice. **a**, UMAP with Cluster-X overlay showing the distribution of the clusters of immune cells identified by multiparametric spectral flow cytometry in the circulation of young (3 months old, mo) and old (24 mo) mice. **b**, Quantification of the NLR in the circulation of young (3 mo) and old (24 mo) mice (n = 7-10 mice per group). **c**, Representantive dot plots and quantification of CXCR4^hi^ CD62Ll° and CXCR4^lo^ CD62L^hi^ neutrophils in the circulation of young (3 mo) middle-aged (15 mo) and old (24 mo) mice (n = 4-23 mice per group). **d**, Schematic diagram depicting the adoptive transfer of blood from 3 mo CD45.1^+^ mice into either young (3 mo) or old (24 mo) CD45.2^+^ recipients. **e**,**f**, Contour plot (**e**) and quantification (**f**) showing CD45.1^+^ CXCR4^hi^ CD62L^lo^ transferred neutrophils before (0 hr) and after (4 hr) the adoptive transfer into either young (3 mo) or old (24 mo) mice (n = 5 mice per group). **g**, Percentage of HSCs, and GMPs in the BM from young (3 mo) and old (24 mo) mice assessed by flow cytometry (n = 6-8 mice per group). **h**, Representative flow cytometry plots and quantification of the percentage of CD4^+^ cells in the BM of young (3 mo) and old (24 mo) mice. **i**, Schematic diagram depicting the adoptive transfer of CD4^+^ cells isolated from young (3 mo) or old (24 mo) into young *Rag1^-/-^* recipients. **j**, Quantification of the percentage of LS^-^K, GMPs and neutrophils in the BM of recipient animals 7 days after the adoptive transfer (n = 6-7 mice per group).

According to previous data, this rise in neutrophils correlated with a skewing of haematopoiesis during ageing (Fig. 1g and Extended Data Fig. 2). Aged mice exhibited increased percentages of haematopoietic stem cells (HSCs) and granulocyte-monocyte precursors (GMPs) (Fig. 1g), together with increased numbers of neutrophils and monocytes in the BM (Extended Data Fig. 1g).

As recent evidence supports the notion that T cells infiltrate the BM and influence haematopoiesis during stressful and autoimmune conditions ^13–16^, we wondered whether this mechanism also operates in old mice. We observed that, unlike in the blood, lymph nodes and spleen, old mice exhibited an increased percentage of CD4^+^ T cells in the BM (Fig. 1h and Extended Data Fig. 3a-c). To directly dissect the capacity of aged CD4^+^ T cells to induce granulopoiesis, we performed adoptive transfer experiments of CD4^+^ T cells from young and old mice into T- and B-cell-deficient recipient (*Rag1^-/-^*) mice ^19^ and we analysed the frequency of BM hematopoietic precursors 7 days post-transfer (Fig. 1i). Compared with the transfer of CD4^+^ T cells from young mice, infusion of CD4^+^ T cells from old mice led to enhanced frequencies of LS^-^K cells and downstream granulocyte-monocyte progenitors (GMPs) together with an increased frequency of BM neutrophils (Fig. 1j), indicating that old CD4^+^ T cells promote granulopoiesis. Further supporting a role of T cells in the regulation of myeloid cell generation, old *Cd3e^-/-^* mice lacking T cells exhibited lower levels of circulating neutrophils and monocytes than age-matched control mice (Extended Data Fig. 1h).

Altogether, these results indicate that T cells accumulate in the BM during ageing leading to enhanced granulopoiesis and neutrophil output.

### CCL5^+^ CD4^+^ CTLs accumulate in the bone marrow during ageing

Single-cell analyses have uncovered the heterogeneity of T cells during ageing in different organs such as the spleen, peritoneum, lung and liver ^20,21^. Based on these observations, we implemented a panel of 15 antibodies that allows to explore the diversity of CD4^+^ T cells in the BM during ageing by spectral flow cytometry (Fig. 2a,b and Extended Data Fig. 4). Unbiased analysis of CD4^+^ T cells from the BM by UMAP and ulterior clusterization identified 9 clusters, including naïve T cells (cluster 1 or “Naïve”), resting and activated T regulatory cells (clusters 2 and 3 or “rTregs” and “aTregs”, respectively). Among the rest of the clusters, we identified T central memory T cells (cluster 4 or “TCM”), effector memory T cells (cluster 5 or “TEM”), PD-1^+^ T cells (cluster 6 or “PD-1^+^”), a cluster defined by the co-expression of CXCR6 and CD38 (cluster 7 or “CXCR6^+^”), a CCL5^+^ cluster (cluster 8 or “CCL5^+^”) and a cluster lacking expression of most of the assessed markers (cluster 9) (Fig. 2a,b and Extended Data Fig. 4). We compared the proportion of all subsets in old versus young mice (Fig. 2c). As expected, the two naïve subsets (clusters 1 and 2) were significantly enriched in young mice and the aTregs cluster showed a similar abundance in both groups of age. Notably, the TCM, TEM and CXCR6^+^ clusters were enlarged in young mice. Conversely, T cells included in the PD-1^+^ cluster increased in old mice. Critically, the vast majority of BM CD4^+^ T cells showing a significant accumulation in old mice grouped in the CCL5^+^ cluster, accounting for >25% of the CD4^+^ cells while it was negligible in young mice [CCL5^+^: mean young/old: 1.64/26.78, p < 0.0001] (Fig. 2c). The CCL5^+^ cluster was characterized by expressing high levels of the activation marker CD38, the transcription factor Eomesodermin (EOMES) and the checkpoint inhibitor PD-1 together with a low expression of CD44 (Fig. 2b and Extended Data Fig. 4), resembling cytotoxic T cells (CD4^+^ CTLs)^20^. Positive expression of Perforin in most BM CD4^+^ CCL5^+^ cells from old mice further confirmed the cytotoxic identity of this cluster (Fig. 2d).

**Figure 2.**
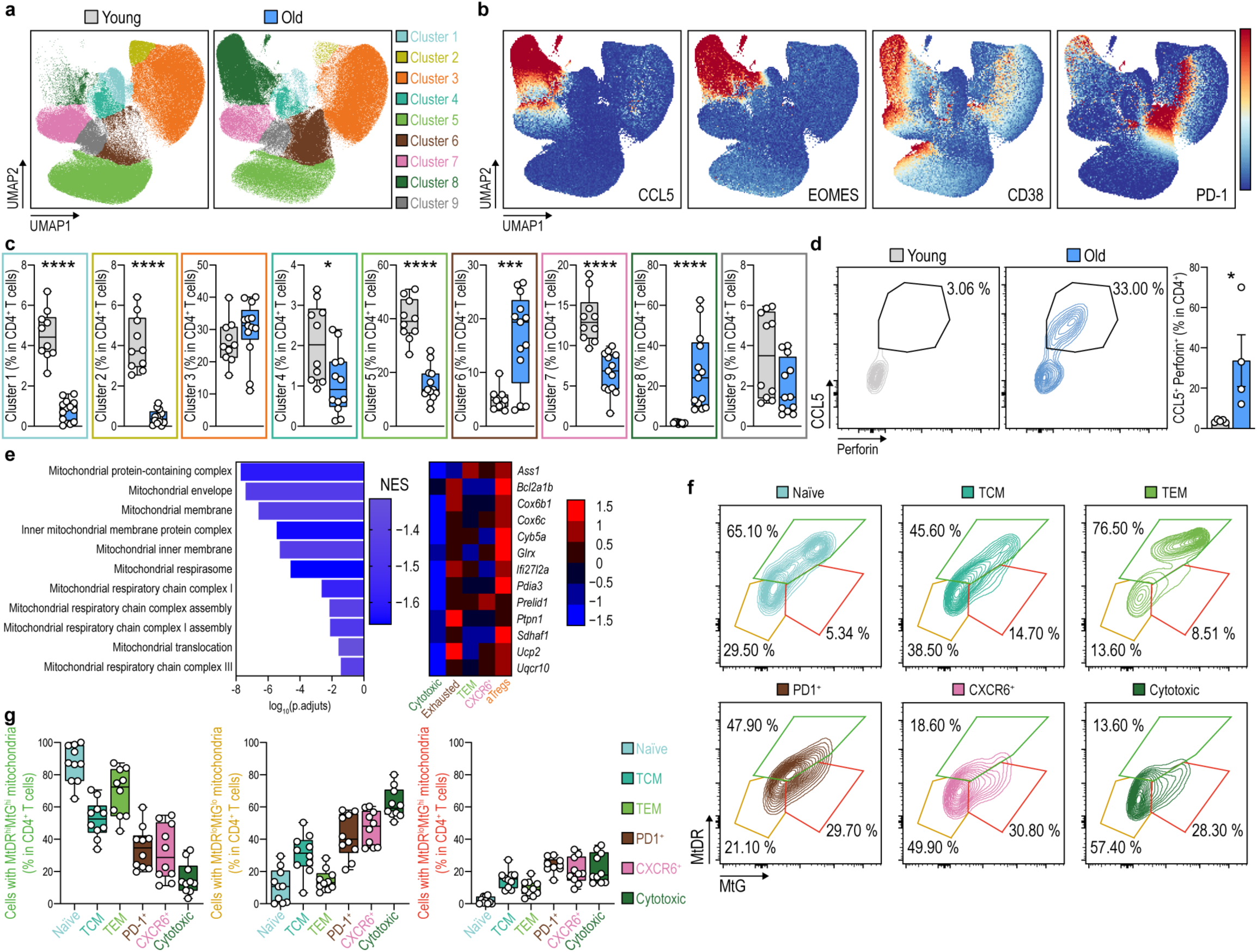
CD4^+^ T cells accumulating in the aged bone marrow exhibit a cytotoxic profile and harbour dysfunctional mitochondria. a, UMAP with Cluster-X overlay showing the distribution of the nine clusters of CD4^+^ T cells identified by multiparametric spectral flow cytometry in the BM from young (3 months old, mo) and old (24 mo) mice. **b**, UMAP representation of the expression levels of representative markers used to identify the CD4^+^ CTLs. **c**, Boxplots comparing the percentage of cells in each T cell subset in young (3 mo) and old (20 mo) mice (n = 10-13 mice per group). **d**, Representative flow cytometry plots and quantification of the percentage of CCL5^+^ Perforin^+^ CD4^+^ cells in the BM of young (3 mo) and old (24 mo) mice (n = 4-5 mice per group). **e**, GSEA analysis of differentially expressed pathways related to mitochondrial function in CD4^+^ CTLs compared with the other differentiated CD4^+^ cells (left) and heat map showing the expression of differentially expressed genes between the different clusters (right). **f**, Representative contour plots showing MtDR and MtG in the clusters identified by extracellular staining in BM CD4^+^ cells from old (24 mo) mice (n = 10 mice per group). **g**, Quantification of the percentage of BM T cells exhibiting healthy (MtDR^hi^MtG^lo^, left), depolarized (MtDR^lo^MtG^lo,^ middle) or (MtDR^lo^MtG^hi^, right) mitochondria in the different clusters.

### Bone marrow CD4^+^ CTLs harbour depolarized mitochondria

T cell ageing correlates with a mitochondrial function decline ^22,23^. We tested whether CD4^+^ CTLs in old mice exhibit mitochondrial dysfunction by using the Mitotracker Green probe (MtG), which stains the whole mitochondrial mass pool, in combination with Mitotracker Deep Red (MtDR), whose incorporation depends on the mitochondrial membrane potential. This technique permits the identification of cells harbouring healthy mitochondria (MtDR^hi^MtG^hi^) or depolarized mitochondria (MtDR^lo^MtG^lo^, and MtDR^lo^MtG^hi^). Comparison of CD4^+^ T cells from the BM, spleen, and Peyer’s patches revealed that MtDR^lo^MtG^lo^ and MtDR^lo^MtG^hi^ CD4^+^ T cells more frequently accumulated in the BM than in the rest of the tissues in old mice (Extended Data Fig. 5a-c), suggesting that aged T cells with dysfunctional mitochondria preferentially accumulate in the BM.

To identify which of the T cell clusters harbour dysfunctional mitochondria, we first performed analysis of publicly available scRNAseq databases applying Gene Set Enrichment Analysis (GSEA) to compare significantly altered pathways between CD4^+^ CTLs and the rest of the CD4^+^ T cell subsets ^20^. This approach evidenced a downregulation of several mitochondrial function-related pathways in the CD4^+^ CTL cluster (Fig. 2e). Hence, we evaluated the mitochondrial membrane potential in the different T cell subsets in the BM. To overcome the impossibility of fixing the MtG dye, we set up a panel of antibodies against extracellular markers to identify the different clusters of BM T cells (Extended Data Fig. 6a). CD4^+^ CTLs were identified as CD44^lo^CD62L^-^CD38^hi^ and intracellular staining for CCL5 confirmed that the majority of CCL5^+^ cells were included in this population (Extended Data Fig. 6b and Supplementary Table 1). With this strategy, we observed similar proportions of the different clusters in the aged BM to that observed by unbiased clusterization, with a remarkable accumulation of CD4^+^ CTLs (Extended Data Fig. 6b,c). Of note, cells identified as CD4^+^ CTL contained the highest percentage of T cells with MtDR^lo^MtG^lo^ and MtDR^lo^MtG^hi^ depolarized mitochondria (Fig. 2f,g and Supplementary Table 2) and therefore showed a concomitantly decreased MtDR/MtG ratio (Extended Data Fig. 6d and Supplementary Table 1). Taken together, these results support the notion that T cells with dysfunctional mitochondria exhibiting a CD4^+^ CTL profile accumulate in the BM during ageing.

### Increased granulopoiesis in mice with mitochondrial dysfunction in T cells

In light of the accumulation of depolarized mitochondria in CD4^+^ CTLs, we investigated whether a deteriorated mitochondrial function in CD4^+^ T cells was sufficient to induce CTL conversion and granulopoiesis skewing. To this end, we used mice with a T cell-specific deletion of the mitochondrial transcription factor A (*Tfam*) gene by crossing Tfam^fl/fl^ mice with mice expressing the CRE recombinase under the *Cd4* promoter (CD4^Cre^) (Extended Data Fig. 7a). T cells from CD4^Cre^ Tfam^fl/fl^ mice display an activated phenotype ^6,24^, inducing the expression of inflammageing-associated cytokines that trigger paracrine senescence and age-associated multimorbidity^6^. At 8 months old, CD4^Cre^ Tfam^fl/fl^ mice presented an accumulation of CD4^+^ CTLs similar to observed in 24 months old mice (Extended Data Fig. 7b, Fig. 2c). Analysis of circulating immune cell populations showed a remarkable accumulation of neutrophils together with a decrease of T and B cells (Extended Data Fig. 7c,d and Extended Fig. 2a) that correlated with a concomitantly increased NLR (Extended Data Fig. 7e), mirroring observations in old mice (Fig. 1a,b and Extended Fig. 1b). Supporting the idea that enhanced granulopoiesis contributes to the increased levels of circulatory neutrophils, we detected an increased percentage of GMPs in the BM of CD4^Cre^ Tfam^fl/fl^ mice (Extended Data Fig. 7f). CD4^Cre^ Tfam^fl/fl^ mice also presented increased levels of circulating pro-inflammatory CXCR4^hi^CD62L^lo^ neutrophils (Extended Data Fig. 7g) and increased numbers of splenic neutrophils and monocytes (Extended Data Fig. 7h) together with infiltrating myeloperoxidase-positive (MPO) neutrophils in the kidney, liver and lung (Extended Data Fig. 7i,j). Altogether, these findings suggest that mitochondrial dysfunction in T cells promotes CD4^+^ CTL differentiation and granulopoiesis induction, mirroring observations in naturally aged mice.

### Prolonged TCR stimulation in an aged milieu induces mitochondrial dysfunction and CCL5^+^ CD4^+^ CTL differentiation

To investigate whether the differentiation of T cells towards CCL5^+^ CD4^+^ CTL is triggered by exposure to an aged environment, we performed heterochronic adoptive transfer of CD45.1 CD4^+^ T cells isolated from young mice into either young or old CD45.2 recipient mice (Fig. 3a). Analysis of T cells 14 days post-transfer showed an increased percentage of CD45.1^+^ CD4^+^ CTLs in the blood of old compared to young recipient mice (Fig. 3b). By 21 days post-transfer, old mice presented ∼ 60% of CD45.1^+^ BM T cells exhibiting a CD4^+^ CTL profile, contrasting with a ∼ 3% in the BM of young recipients (Fig. 3c). Additionally, ageing of the host correlated with an increased infiltration of CD45.1 CD4^+^ CTLs in different tissues. Of note, comparison of the percentage of CD45.1 CD4^+^ CTL in the BM, the liver, the white adipose tissue and the colon, revealed that donor cells were particularly enriched in the BM during ageing, with a remarkable 18.5-fold increase of young levels, confirming that the BM is a preferential site for CD4^+^ CTL migration (Fig. 3d).

**Figure 3.**
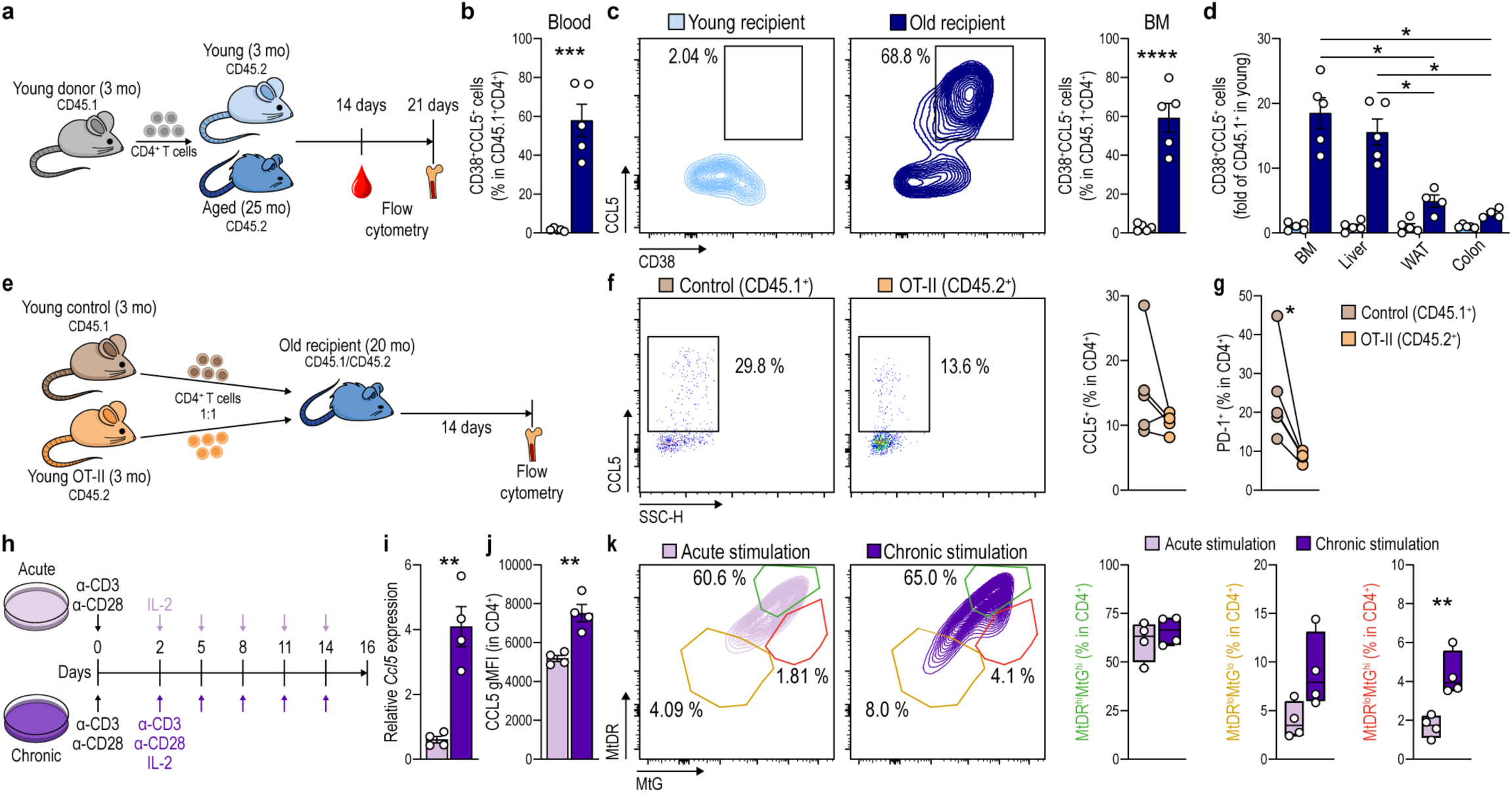
Prolonged TCR engagement in an aged environment induces mitochondrial dysfunction and CD4^+^ CTL differentiation. **a**, Schematic diagram depicting the adoptive transfer of CD45.1^+^ T cells isolated from 3 months old (mo) mice into either young (3 mo) or old (24 mo) CD45.2^+^ recipients. **b**, Percentage of blood CD4^+^CD38^+^CCL5^+^ T cells in young or old recipients 21 days after the adoptive transfer of CD4^+^ T cells (n = 5 mice per group). **c**, Representative contour plots and quantification of the percentage of BM CD4^+^CD38^+^CCL5^+^ T cells in young or old recipients 21 days after the adoptive transfer of CD4^+^ T cells (n = 5 mice per group). **d**, Fold increase of CD4^+^CD38^+^CCL5^+^ T cells in the indicated organs from young and old recipients (n = 5 mice per group). **e**, Schematic diagram depicting the mixed adoptive transfer of CD45.1^+^ control and CD45.2^+^ OT-II derived cells into old (20 mo) recipients. **f**, Representative dot plots and quantification of the percentage of BM CCL5^+^ CD4^+^ cells in either CD45.1^+^ control or CD45.2^+^ OT-II 14 days after the adoptive transfer (n = 5 mice per group). **g**, Percentage of CD4^+^ PD-1^+^ cells in CD45.1^+^ control or CD45.2^+^ OT-II transferred cells (n = 5 mice per group). **h**, Schematic diagram depicting the protocol followed to acutely and chronically stimulate isolated CD4^+^ T cells in culture by exposure to anti-CD3/anti-CD28 and IL-2. **i**,**j**, Levels of *Ccl5* mRNA (**i**) and CCL5 protein (**j**) in acutely and chronically stimulated T cells determined by RT-qPCR analysis and flow cytometry, respectively (n = 4). **k**, Representative contour plots showing MtDR and MtG and quantification of the percentage of CD4^+^ cells exhibiting healthy (MtDR^hi^MtG^lo^) or depolarized (MtDR^lo^MtG^lo^ or MtDR^lo^MtG^hi^) mitochondria in acutely stimulated versus chronically stimulated conditions (n = 4).

To assess if the conversion to CCL5^+^ CD4^+^ CTL in an aged environment was due to chronic TCR stimulation or to exposure to inflammageing-associated cytokines, we adoptively transferred CD4^+^ cells isolated from young CD45.1 and OT-II CD45.2 mice into old CD45.1.2 recipients in a 1:1 ratio and we analysed the differential infiltration of donor CD4^+^ cells to the BM 14 days later (Fig. 3e). CD4^+^ cells from OT-II mice are genetically engineered to express a TCR that exclusively recognizes a specific peptide of the chicken ovalbumin. In the absence of this peptide, OT-II CD4^+^ cells cannot engage TCR-dependent activation while remaining susceptible to cytokine-mediated activation^25^. We detected a decreased percentage of both CCL5^+^ and PD-1^+^ OT-II CD45.2 T cells in the BM of recipient mice 14 days after the adoptive transfer (Fig. 3f,g), suggesting that TCR signalling is required for CTL differentiation in an aged environment. Furthermore, we compared CD4^+^ cells acutely and chronically (16 days) stimulated in culture with anti-CD3/anti-CD28 (Fig. 3h) and we observed an increased expression of CCL5 (Fig. 3i,j), together with CD38, PD-1 and EOMES (Extended Data Fig. 8a-c), in chronically relative to acutely TCR-stimulated cells. Furthermore, we observed that continuous TCR stimulation led to the accumulation of dysfunctional mitochondria (MtDR^lo^MtG^lo^ and MtDR^lo^MtG^hi^) in cultured CD4^+^ T cells (Fig. 3k). In sum, these results support the notion that exposure to an aged environment induces the differentiation of CD4^+^ cells into CD4^+^ CTLs, requiring prolonged TCR stimulation.

### Bone Marrow CD4^+^ CTLs induce granulopoiesis by CCL5/CCR5 signalling

Given that signalling through CCL5 and CCR5 has been involved in the induction of myeloid skewing in mouse models of multiple sclerosis ^15,26^, we studied the contribution of this pathway to T cell-induced granulopoiesis during ageing. Notably, we initially observed that HSCs from old mice exhibit an upregulation of CCR5, the main CCL5 receptor, leading to increased percentages of CCR5^+^ HSCs in the BM of old mice (Fig. 4a). To explore the relevance of signalling through CCL5/CCR5 in T cell-induced granulopoiesis during ageing, we generated mixed BM chimeric mice by injecting 50% of CD45.1 CCR5^+/+^ and 50% of CD45.2 CCR5^-/-^ BM cells into irradiated CD45.1.2 recipients. Two months later, we adoptively transferred CD4^+^ cells isolated from aged mice and studied the effect in HSCs and downstream precursors of both genotypes one week later (Fig. 4b). Flow cytometry analysis revealed a decreased percentage of LS^-^K cells and GMPs in CD45.2 CCR5^-/-^ cells (Fig. 4c and Extended Fig. 2) that correlated with decreased proportions of CD45.2 CCR5^-/-^ neutrophils and Ly6C^hi^ monocytes in the BM (Fig. 4d). We also confirmed reductions in the percentages of blood CD45.2 CCR5^-/-^ neutrophils and Ly6C^hi^ monocytes (Fig. 4e). Supporting a selective effect on the myeloid lineage, we did not find reductions in the levels of either BM CD45.2 CCR5^-/-^ CD8^+^ cells (Fig. 4f) or blood CD45.2 CCR5^-/-^ CD8^+^ cells (Fig. 4g). In aggregate, these findings suggest that CCR5 signalling in HSCs is involved in T cell-induced granulopoiesis in aged mice.

**Figure 4.**
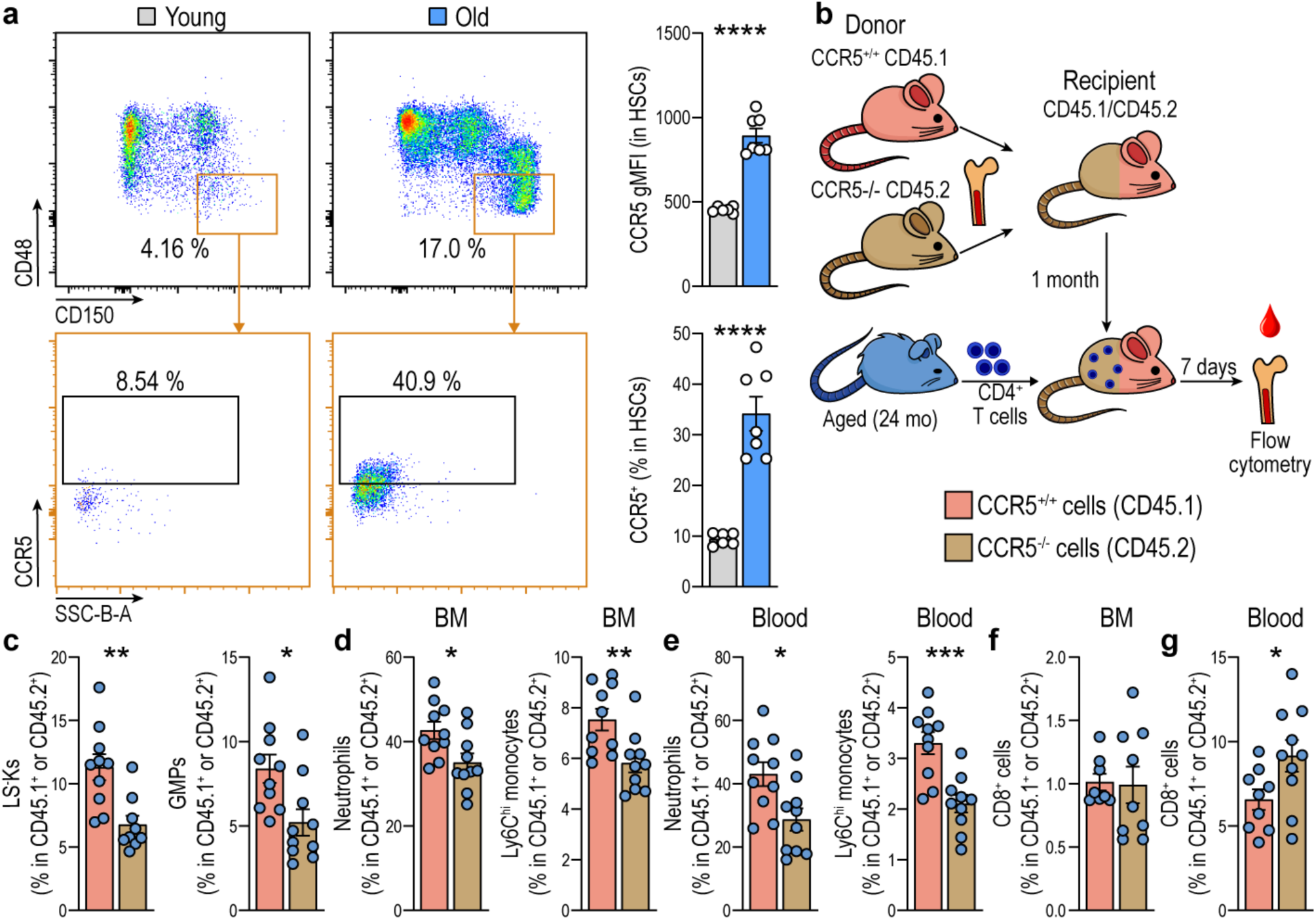
T cells from aged mice induce granulopoiesis via CCL5/CCR5. **a**, Representative dot plots showing the identification of CCR5^+^ HSCs and quantification of the levels of CCR5 and the percentage of CCR5^+^ HSCs in the BM of young (3 months old, mo) and old (24 mo) mice (n = 5-7 per group). **b**, Schematic diagram depicting the strategy followed to generate mixed BM chimeric mice. CCR5^+/+^ (CD45.1^+^) and CCR5^-/-^ (CD45.2^+^) derived BM cells were mixed 1:1 and i.v. injected into lethally irradiated young (5-6 mo) CD45.1.2^+^ recipients. One month after transplantation, CD4^+^ CD45.2^+^ isolated from old (24 mo) mice were adoptively transferred into the mixed chimeric recipients. **c,d,** Quantification of the percentage of LS^-^K and GMPs (**c**), neutrophils and Ly6C^hi^ monocytes (**d**) in CCR5^+/+^ CD45.1^+^ or CCR5^-/-^ CD45.2^+^ cells into the BM of mixed chimeric recipients (n = 10 mice per group). **e**, Quantification of the percentage of circulating neutrophils and Ly6C^hi^ monocytes in mixed chimeric recipients (n = 10 mice per group). **f,g** Percentage of CD8^+^ cells in the BM (**f**) and the blood (**g**) of mixed chimeric recipients (n = 10 mice per group).

### Inhibition of CCR5 reverts myeloid skewing, decreases the NLR and improves health status in aged mice

To investigate whether interfering with CCR5 signalling could prevent age-associated changes in haematopoiesis, we treated old mice for one month with Maraviroc, an FDA-approved CCR5 antagonist ^27^ (Fig. 5a). Treatment with Maraviroc decreased the percentage of GMPs and LS^-^K cells while increased the frequencies of common lymphoid progenitors (CLPs) in old mice (Fig. 5b and Extended Fig. 2), confirming a reversion of the skewing of haematopoiesis. Notably, treatment with Maraviroc decreased the percentage of circulating neutrophils (Fig. 5c) and increased the percentage of CD4^+^ and CD8^+^ T cells (Fig. 5d), leading to a concomitant normalisation of the NLR (Fig. 5e) in old mice. These results suggest that CCR5 inhibition turns haematopoiesis into a more youthful state.

**Figure 5.**
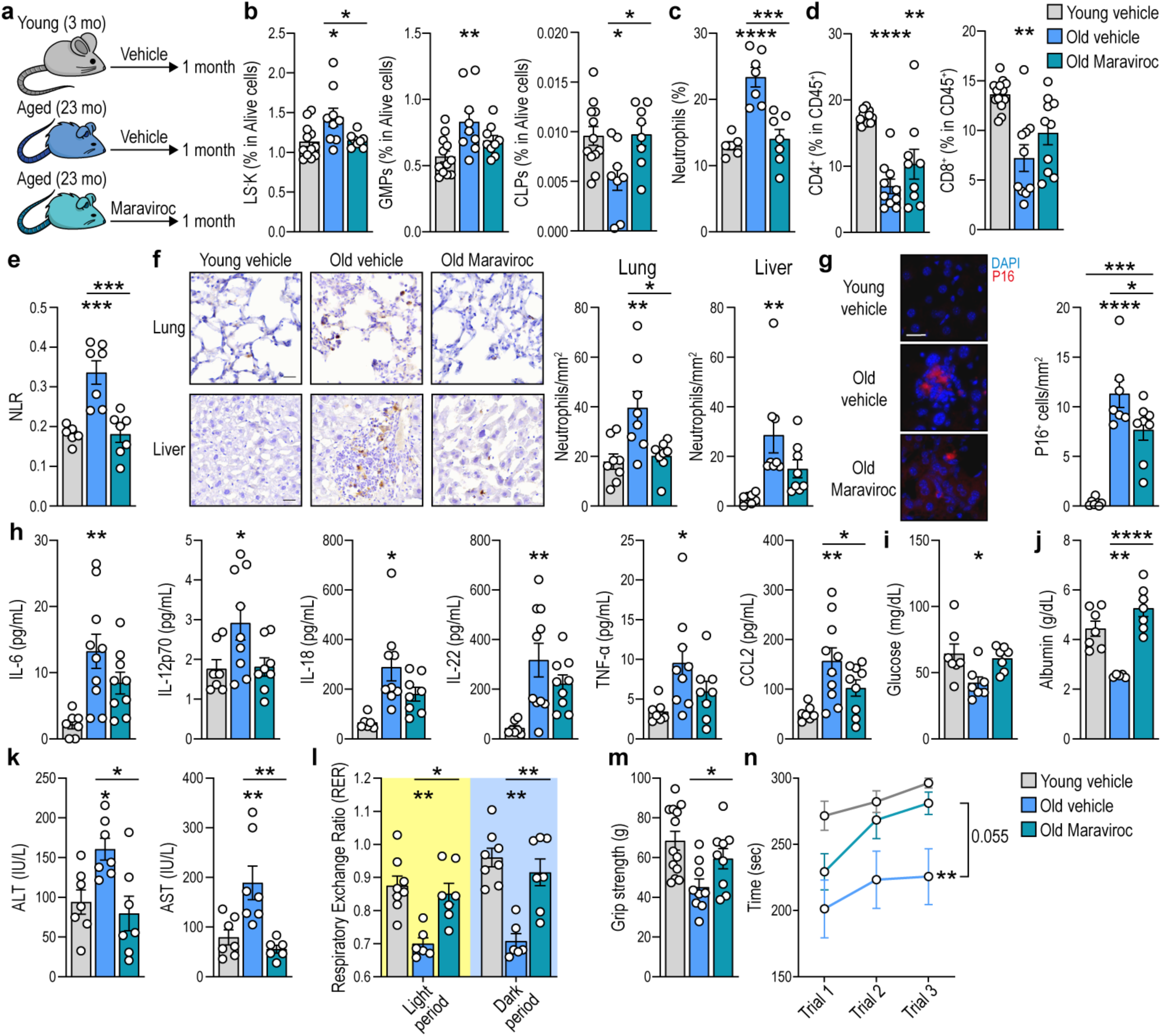
Pharmacological inhibition of CCR5 restores granulopoiesis and improves health status in aged mice. **a**, Schematic diagram detailing the treatment with Maraviroc. Old mice were either daily treated with vehicle or Maraviroc for one month. Young mice were treated in parallel. **b**, Percentage of GMPs, LS^-^K and CLPs in the BM of the three experimental groups (n = 9-12 mice per group). **c**, Percentage of blood neutrophils in the experimental groups measured by blood haematology (n = 6-7 mice per group). **d**, Percentage CD4^+^ and CD8^+^ cells in CD45^+^ cells measured by flow cytometry in blood samples from the experimental groups (n = 9-12 mice per group). **e**, NLR in blood from the experimental groups (n = 6-7 mice per group). **f**, MPO immunohistochemistry and quantification of MPO^+^ cells per mm^2^ in liver and lung sections from all experimental groups (n = 8 mice per group). **g**, Immunofluorescence staining for p16 and quantification of p16^+^ cells in liver sections from the experimental groups (n = 8 mice per group). Scale bar, 20 μm. **h**, Serum levels of the indicated cytokines and chemokines detected by multiplex in the three experimental groups (n = 7 to 9 mice per group). **i**, Serum glucose levels in non-fasted mice from the experimental groups (n = 7 mice per group). **j**, Serum albumin levels in non-fasted mice from the experimental groups (n = 7 mice per group). **k**, Serum ALT and AST levels in non-fasted mice from the experimental groups (n = 7 mice per group). **l**, Respiratory exchange ratio (VCO_2_/VO_2_) assessed in metabolic cages during the light and dark phases among the specified experimental groups (n = 6-8 mice per group). **m**, Quantification of forelimbs strength in the three experimental groups (n = 9-13 mice per group). **n**, Quantification of the latency to fall in the rotarod test in the three experimental groups (n = 6-10 mice per group).

We next studied whether restoring haematopoiesis with Maraviroc improves health outcomes in old mice. Old mice treated with Maraviroc exhibited reduced levels of infiltrating neutrophils in lung and liver (Fig. 5f), which correlated with a decreased presence of senescent cells, identified as p16^+^, in the liver (Fig. 5g). Furthermore, Maraviroc lowered the levels in serum of multiple inflammageing-associated cytokines (Fig. 5h), non-fasting glucose (Fig. 5i), albumin (Fig. 5j), alanine aminotransferase (ALT) and aspartate aminotransferase (AST) (Fig. 5k). Notably, this correlated with the normalisation of different physiological parameters affected in old mice including metabolic alterations such as the respiratory exchange ratio (Fig. 5l), the loss of muscle strength (Fig. 5m) and the impaired locomotor coordination (Fig. 5n). Taken together, these results suggest that CCR5 inhibition by Maraviroc normalises granulopoiesis, leading to improved health outcomes in old mice.

## Discussion

CD4 CTLs are emerging as age-associated T cells, as their numbers correlate with ageing ^20,28^. However, their origin and contribution to ageing are still debated. Previous findings reported the presence of these cells mainly in conditions characterized by continuous TCR stimulation such as viral infections ^29^. We found that young T cells exposed to an aged environment differentiate into CD4^+^ CTL, requiring prolonged TCR activation. It is tempting to speculate that an altered repertoire of presented antigens could be the origin of CD4^+^ CTL differentiation in an aged environment. Regarding their function, CD4^+^ CTLs are responsible of eliminating senescent cells in the skin ^30^ and may be involved in the control of tumour growth ^31^. Despite these potentially beneficial functions, a massive accumulation of CD4^+^ CTL in the BM during ageing, either due to enhanced migration or to delayed clearance, would result in a detrimental increase in granulopoiesis and neutrophilia.

A skewing of haematopoiesis towards the production of myeloid cells is an ageing feature with a major impact on immune system function, tissue homeostasis and healthspan. Even though a high NLR is associated with bad prognoses in most age-associated diseases ^2–5^, the mechanisms driving altered haematopoiesis during ageing remain incompletely understood. Previous findings have established that remodelling of the BM stroma induces the skewing of haematopoiesis in aged mice. Accordingly, inflammation triggered by β2/β3 adrenergic receptors induces haematopoiesis skewing ^12,32^, at least in part, by increasing IL-1β levels in aged mice ^33^. Similarly, decreased levels of trophic factors such as IGF-1 ^34^ or an altered cellular composition of the BM niche, including a Notch-dependent reduction of blood vessels ^35^ or an accumulation of adipocytes ^36^, alter the fate of haematopoietic precursors. Our findings suggest that infiltration of CD4^+^ CTLs into the BM during ageing will be a critical factor contributing to the remodelling of the BM niche. These cells, characterized by the overexpression of CCL5, induce granulopoiesis by signalling to CCR5 in haematopoietic precursors. These observations are in line with previous findings reporting enhanced granulopoiesis and neutrophil generation by BM infiltrating autoreactive T cells in multiple sclerosis ^15^.

We found enhanced granulopoiesis resulting in elevated pro-inflammatory CXCR4^hi^CD62L^lo^ neutrophils in old mice. Remarkably, recent evidence suggest that pro-inflammatory neutrophils can induce paracrine senescence ^11^, trigger age-related lung damage ^10^, worsen myocardial infarction ^37^ and foster skin inflammation ^38^. Importantly, we found that enhanced granulopoiesis in aged mice is prevented by pharmacological inhibition of CCR5, correlating with an improvement of multiple ageing-associated conditions. Supporting these findings, human individuals carrying homozygous loss of function mutations in CCR5 (CCR5-Delta32) present better outcomes after stroke ^39^ and during multiple sclerosis ^26^. Future studies should investigate whether this improvement is mediated by a decreased induction of granulopoiesis, which contributes to aggravating stroke and autoimmune neuroinflammation in mice ^15,40^.

In all, these findings place T cells as inducers of the disbalance in haematopoiesis seen during ageing. As a high NLR is a predictor of mortality in most age-related diseases ^2–5^, and Maraviroc is already an FDA-approved drug for the treatment of HIV patients ^27^, our results may have broad therapeutic implications for the treatment of age-related pathologies. These findings set the stage for the development of drugs to eliminate CD4^+^ CTLs, avoid their recruitment into the BM, and modify their interplay with haematopoietic precursors to improve age-associated outcomes.

## Acknowledgements

The authors thank Maria Navarro and César Cobaleda for critical discussion of the manuscript. Research in the Mittelbrunn lab was supported by European Research Council (ERC-2021-CoG 101044248-Let T Be), by Comunidad de Madrid (Spain) (Y2020/BIO-6350 NutriSION-CM synergy) and by Spanish Ministerio de Ciencia e Innovación (PID2022-141169OB-I00) grants. G.S-H. was supported by an FPI-UAM grant from the Universidad Autónoma de Madrid and by Let T Be. E.G-R and I.F-Q were supported by a Juan de la Cierva-Incorporación grant (IJC2018-036850-I and JC2020-044392, respectively) from the Ministerio de Ciencia, Innovación y Universidades (Spain) and by ERC-2021-CoG 101044248-Let T Be. M.M.G.H. and J.I.E.-L. were supported by FPU grants (FPU19/02576 and FPU20/04066, respectively), both from Ministerio de Ciencia, Innovación y Universidades (Spain). C.A., A.G., R.dC., and M.G. are supported by the NIA IRP, NIH. We thank the Mouse Metabolism & Phenotyping core in Baylor College of Medicine at Houston, Texas (USA) for analysis of serum samples and the flow cytometry facility of CBMSO.

## Methods

### Animal procedures

All mice were bred and aged in the specific-pathogen-free facility of Centro de Biología Molecular Severo Ochoa (CBMSO, Madrid, Spain) in accordance with European Union recommendations and institutional guidelines. C57BL/6J HccRsd mice were purchased from Envigo or bred at the CBMSO animal facilities. *Tfam*^fl/fl^ mice were kindly provided by N.G. Larsson ^41^, and CD4^Cre+/wt^ mice were purchased from the Jackson Laboratory. Double heterozygotes (*Tfam*^+/fl^, CD4^Cre+/wt^) were obtained and backcrossed to the *Tfam*^fl/fl^ strain to generate cell-specific knockouts (CD4^Cre^ *Tfam*^fl/fl^). OT-II mice were kindly provided by Dr. N. Martínez-Martín (CBMSO). CCR5^-/-^ mice were provided by Prof. S. Mañes (CNB). *Rag1^-/-^* (1547488) ^19^, C57BL/6J CD45.1.2 and C57BL/6J CD45.1 mice were provided by Dr. C. Cobaleda (CBMSO). Surgical and experimental procedures were approved by the ethics committee of the Consejo Superior de Investigaciones Científicas (CSIC). Both male and female mice were used in this study. In general, mice were used at different ages: young (less than 4 mo), old (22-25 mo) unless otherwise specified in the figure legend.

#### Adoptive transfer of immune cells

For neutrophil analysis, whole blood from four donor C57BL/6 CD45.1 mice, previously euthanized by exposure to CO_2_, was isolated by cardiac puncture. Red blood cells were removed in erythrocyte lysis buffer for 7 min and immune cells were resuspended in 0.9% NaCl. Equal amounts of immune cells were i.v. injected into young or old CD45.2 recipients. The percentage of CXCR4^hi^CD62L^lo^ and CXCR4^lo^CD62L^hi^ was analyzed in Ly6C^+^ Ly6G^+^ neutrophils before and four hours after the adoptive transfer.

For the adoptive transfer of CD4^+^ cells, the spleen and the axillary, mesenteric and inguinal lymph nodes were harvested from either young and old C57BL/6 CD45.1, C57BL/6 CD45.2 or OT-II (CD45.2) donor mice as specified in figures. Organs were mashed and filtered through a 70-µm cell strainer and erythrocytes were removed in erythrocyte lysis buffer for 5 min. CD4^+^ cells were purified with the MojoSort^TM^ mouse CD4 T cell isolation kit (Biolegend, 480006) according to manufacturer instructions. CD4^+^ cells were resuspended in 0.9% NaCl and 6 million cells were i.v. injected into every young and old C57BL/6 CD45.1, CD45.2 or CD45.1.2 recipient as specified in the figure legends. The purity of CD4^+^ isolated cells was routinely tested by flow cytometry, typically ranging from 85-95% of total cells.

#### BM transplantation

For BM transplantation, recipient mice were lethally irradiated with two doses of 5.5 Gy and intravenously injected with 5 x 10^6^ cells. Mice were administered with 1 mg/ml of neomycin in the drinking water for four weeks starting one week before transplantation. Recipient mice were allowed to reconstitute immune cell populations for two months and reconstitution was routinely assessed before any further intervention by flow cytometry.

#### Treatment with Maraviroc

Maraviroc (MedChem Express) was dissolved in 10% DMSO (Thermo Scientific, 20688), 40% PEG300 (Selleckchem, S6704), 5% Tween-80 (Selleckchem, S6702) and 45% saline and the resultant solution was sonicated. Young (3 mo) and Old (23 mo) C57BL/6J HccRsd mice were assigned in three groups; young mice were i.p. daily treated with vehicle and old mice were i.p. daily treated with either vehicle or 35 mg/kg Maraviroc for one month.

#### Rotarod test

Motor coordination was assessed by performing the rotarod test in an accelerating rotarod apparatus (Ugo Basile, Varese, Italy). Mice were trained for 1 day at a constant speed: two times at 4 r.p.m. for 1 min and two times with acceleration from 4-8 r.p.m. for 1 min. On the second day, the rotarod was set to progressively accelerate from 4 to 40 r.p.m for 5 min. Mice were tested three times. During the accelerating trials mice were video-recorded and the latency to fall from the rod was measured.

#### Forelimbs strength analysis

Limb grip strength was measured as tension force using a digital force transducer (Grip Strength Meter, Bioseb). Ten measurements were performed for each animal, with a 10 second resting period between measurements and average of all measurements of every mouse was compared.

#### Metabolic cages

Metabolic parameters were measured using PhenoMaster TSE-Systems (Germany). Mice were singly housed and acclimatized for 1 days before data monitoring. All parameters were measured continuously and simultaneously for 1 day.

### Tissue processing for flow cytometry

Mice were euthanized with CO_2_ followed by transcardiac perfusion with ice-cold PBS. The indicated tissues were extracted and processed as specified:

#### Spleen

Spleen was mashed and filtered through a 70-µm cell strainer. Red blood cells were removed in Erythrocyte lysis buffer (ammonium chloride 0.15 M, sodium bicarbonate 0.01 M, EDTA 0.1 mM) for 5 min.

#### Blood

Blood was extracted either from the facial vein or the heart in living or euthanized mice, respectively. Red blood cells were removed in Erythrocyte lysis buffer for 7 min. Cells were washed and stained.

#### Peyer’s Patches or lymph nodes

Peyer’s patches and lymph nodes were harvested from the intestine and mashed into a 70-µm cell strainer. Cell suspension was centrifuged at 400 g for 5 min at 4°C. Cell pellets were resuspended in 2% FBS RPMI.

#### White adipose tissue

Gonadal white adipose tissue was digested into pre-warmed 2 mg/ml BSA 2% FBS RPMI supplemented with 2 mg/ml collagenase type II (Sigma, C6885) and placed under shaking at 180 rpm for 40 min at 37°C. Digested tissues were vertically rested to separate fat from the aqueous phases, which were obtained using a 18G syringe needle. Cell suspensions were filtered through a 70-µm cell strainer and washed with 2% FBS RPMI. Erythrocytes were removed in lysis buffer for 5 min at 4°C and washed with 1 mM EDTA PBS.

#### Liver

Liver was harvested and cut into pre-warmed 25 mM Hepes 10% FBS RPMI supplemented with 0.4 mg/ml collagenase type VIII (Sigma, C2139) under shaking at 180 rpm for 45 min at 37°C. Digested tissue was filtered through a 70-µm cell strainer and centrifuged at 350 g for 5 min at 4 °C. Red blood cells were removed in erythrocyte lysis buffer for 5 min. For leucocyte enrichment, supernatants were centrifuged in a 40%/70% Percoll gradient (Sigma, GE17-0891-01) at 1250 g for 30 min at RT with acceleration on 6 and without brake. Isolated cells were washed with PBS and resuspended in 2% FBS RPMI.

#### Bone marrow

Femurs and tibias were collected. Cells from the bone marrow were obtained by centrifuging the bones at 6000 g for 1 min. Red blood cells were removed in Erythrocyte lysis buffer for 3 min.

### Flow cytometry

To differentiate between live and dead cells were firstly stained with the Zombie NIR™ Fixable Viability Kit (BioLegend, 423106), the Zombie Yellow™ Fixable Viability Kit (BioLegend, 423104) or the Ghost Dye™ Violet 540 (Tonbo Biosciences, 13-0879) for 20 min at 4°C. Cells were washed with FACS staining buffer (PBS supplemented with 2% fetal bovine serum and 1 mM EDTA) and incubated with Fc receptor blocker purified rat anti-mouse anti-CD16/CD32 (BD Biosciences, 553142) for 20 min at 4°C. Cells were then incubated with primary antibodies for 20 min at 4°C and were washed twice with a FACS staining buffer. The following antibodies were used for surface antigen staining:

| Antigen | Fluorochrome | Clone/#Catalog | Supplier | Dilution |
| --- | --- | --- | --- | --- |
| CD8 | BUV615 | 53-6.7 | ThermoFisher | 1:500 |
| CD127 (IL-7R) | BUV737 | SB/199 | BD Horizon | 1:200 |
| Ly6G | BV421 | 1A8 | Biolegend | 1:400 |
| CD186 (CXCR6) | BV421 | SA051D1 | Biolegend | 1:100 |
| Streptavidin | BV421 | 405225 | Biolegend | 1:500 |
| CD38 | Pacific Blue | 90 | Biolegend | 1:500 |
| TIM3 | BV480 | 5D12/TIM-3 | BD Biosciences | 1:200 |
| CD45.2 | V500 | 104 | Tonbo Biosciences | 1:400 |
| CD16/32 | BV510 | 93 | Biolegend | 1:100 |
| CD44 | BV570 | IM7 | Biolegend | 1:200 |
| CD66a (CEACAM1a) | BV650 | MAB-CC1 | Biolegend | 1:200 |
| CD223 (LAG3) | BV650 | C9B7W | Biolegend | 1:100 |
| CD62L | BV711 | MEL-14 | Biolegend | 1:400 |
| Ly6C | BV785 | HK1.4 | Biolegend | 1:400 |
| B220 | FITC | RA3-6B2 | Tonbo Biosciences | 1:200 |
| CD34 | FITC | RAM34 | BD Biosciences | 1:100 |
| CD62L | FITC | MEL-14 | Tonbo Biosciences | 1:400 |
| FoxP3 | FITC | Fjk-16s | Invitrogen | 1:400 |
| Ly6C | PerCP Cy5.5 | HK1.4 | Biolegend | 1:400 |
| CD44 | PerCP Cy5.5 | IM7 | Tonbo Biosciences | 1:200 |
| CD45.2 | PerCP Cy5.5 | 104 | Biolegend | 1:400 |
| CD195 (CCR5) | PerCP Cy5.5 | HM-CCR5 | Biolegend | 1:100 |
| PD1 | PerCP eFluor710 | J43 | ThermoFisher | 1:200 |
| CD135 (FLT3) | PerCP eFluor710 | A2F10 | ThermoScientific | 1:200 |
| CD8 | PE | 53-6.7 | Tonbo Biosciences | 1:400 |
| CD117 (cKit) | PE | ACK2 | Tonbo Biosciences | 1:200 |
| CD184 (CXCR4) | PE | L276F12 | Biolegend | 1:100 |
| CD182 (CXCR2) | PE-Dazzle 594 | SA045E1 | Biolegend | 1:200 |
| NKG2D | PE-Dazzle594 | CX5 | Biolegend | 1:200 |
| CD25 | PE Cy5 | PC61 | Biolegend | 1:400 |
| Ly6G | PE Cy7 | 1A8 | Biolegend | 1:400 |
| CD101 | PE Cy7 | Moushi101 | ThermoScientific | 1:200 |
| SLAM (CD150) | PE Cy7 | TC15-12F12.2 | Biolegend | 1:200 |
| CD54 (ICAM-1) | PE Cy7 | YN1/1.7.4 | Biolegend | 1:200 |
| CX3CR1 | APC | SA011F11 | Biolegend | 1:200 |
| CD11b | APC | M1/70 | Tonbo Biosciences | 1:200 |
| CD284 (TLR4) | APC | SA15-21 | Biolegend | 1:100 |
| CD48 | APC | HM48-1 | Biolegend | 1:200 |
| CD11b | Red-Fluor 710 | M1/70 | Tonbo Biosciences | 1:200 |
| Sca-1 | APC-Cy7 | D7 | Biolegend | 1:200 |
| CD45 | APC-Cy7 | 30-F11 | Biolegend | 1:400 |
| CD45.1 | APC-Cy7 | A20 | Tonbo Biosciences | 1:400 |
| CD279 (PD-1) | APC-Fire750 | 29F.1A12 | Biolegend | 1:400 |
| CD19 | APC-eFluor780 | eBio1D3 | ThermoScientific | 1:200 |
| CD4 | APC Fire810 | Gk1.5 | Biolegend | 1:1000 |
| B220 | Biotin | RA3-6B2 | Tonbo Biosciences | 1:200 |
| Ter119 | Biotin | TER-119 | Tonbo Biosciences | 1:200 |
| Gr-1 | Biotin | RB6-8C5 | Tonbo Biosciences | 1:200 |
| CD11b | Biotin | M1/70 | Tonbo Biosciences | 1:200 |
| CD11c | Biotin | M418 | Tonbo Biosciences | 1:200 |
| CD4 | Biotin | GK1.5 | Tonbo Biosciences | 1:200 |
| CD8 | Biotin | 53-6.7 | Tonbo Biosciences | 1:200 |
| CD3 | Biotin | 145-2C11 | Tonbo Biosciences | 1:200 |

For intracellular staining, cells were fixed and permeabilized using the FoxP3/Transcription Factor Staining Kit (eBioscience) for 20 min at RT in darkness. Cells were then stained with the following intracellular antibodies:

| Antigen | Fluorochrome | Clone/#Catalog | Supplier | Dilution |
| --- | --- | --- | --- | --- |
| FoxP3 | FITC | Fjk-16s | Invitrogen | 1:150 |
| CCL5 | PE | 2E9/CCL5 | Biolegend | 1:200 |
| Eomes | PE-Cy7 | Dan11mag | Invitrogen | 1:200 |
| Perforin | APC | S16009A | Biolegend | 1:200 |
| TOX | eFluor660 | TRX10 | Invitrogen | 1:300 |
| BHLHE40 | Alexa Fluor 700 | #NB100-1800AF700 | Novus Biologicals | 1:200 |

Flow cytometry experiments were performed with either a 4-laser or a 5-laser (Violet, blue, yellow-green, red) Aurora flow cytometer (Cytek Biosciences). Data were analyzed with the FlowJo v10.5.3 software (BD Biosciences). Gating strategies were set on the basis of fluorescence minus one controls and unstained samples. All the samples in the experiment excluded dead cells, clumps, and debris.

#### Analysis of mitochondrial membrane potential

Analysis of mitochondrial mass and mitochondrial membrane potential was performed by flow cytometry in cells labelled for 30 min with 50 nM MitoTracker^TM^ Green FM (Invitrogen, M7514) and 25 nM Mitotracker^TM^ DeepRed FM (Invitrogen, M22426) in RPMI medium in a 37°C and 5% CO_2_ incubator prior to extracellular staining.

### Dimensional reduction and clustering analysis of flow cytometry data

Dimensional reduction and clustering analysis of flow cytometry data was performed using OMIQ (Dotmatics). First, non-lymphocyte cells, doublets and dead cells were excluded based on viability staining and FSC/SSC parameters. Then, 35,000 CD4^+^ cells from each sample were subsampled for further analysis. For dimensional reduction, the Uniform Manifold Approximation and Projection (UMAP) algorithm was applied with the following parameters: Neighbors = 15, Minimum Distance = 0.4, Components = 2, Learning Rate = 1, Epochs = 200. For unbiased clustering, the Cluster-X algorithm was applied with Alpha = 0.001. The mean fluorescence intensity of each marker was projected on the UMAP plots and used to infer the identity of the clusters. Similar clusters were combined.

### Haematology

The ADVIA® 2120i Hematology System (Siemens Healthineers, Erlangen, Germany) was used to quantitatively measure blood variables. For ADVIA measurements, 50 µL of whole blood was resuspended in 150 µL of RPMI-EDTA-K2 medium. The instrument was calibrated on the day of testing as per the manufacturer’s instructions.

### Immunohistochemistry and immunofluorescence

Organs obtained from intracardiac perfused animals were fixed in 10% neutral buffered formalin for 48 hr and dehydrated in 70% ethanol until processing. Dehydrated organs were embedded in paraffin sections and processed with a microtome to generate x μm sections. For immunostaining, the deparaffinized sections were rehydrated and boiled in order to retrieve antigens (10 mM citrate buffer, 0.05% Triton X-100, pH 6). Then, sections were blocked for 45 min in 10% goat serum, 5% horse serum, 0.05% TritonX-100, and 2% BSA in PBS. Endogenous peroxidase and biotin were blocked with 1% hydrogen peroxide-methanol for 10 min and a biotin blocking kit (Vector Laboratories), respectively. Sections were incubated with a goat polyclonal anti-MPO antibody (AF3667, RD Systems) and color was developed with 3,3’-diaminobenzidine (DAB, Vector Laboratories). Sections were counterstained with hematoxylin and mounted in DPX (Fluka). Slides were scanned with a NanoZoomer-RS scanner (Hamamatsu). Briefly, for each animal, the number of MPO-positive cells was quantified in digitized images in which an area covering at least one-half of a section of the whole organ was previously defined. Then, the density of cells per area was estimated by using the Zen 2.3 software (Carl Zeiss, Inc.).

For immunofluorescence, after blocking, sections were instead incubated with a primary mouse monoclonal anti-p16 antibody (ab54210, Abcam) followed by a Donkey anti-mouse fluorophore-coupled secondary antibody (ThermoFisher).

### Analysis of mouse serum samples

All the plasma analysis was done by Mouse Metabolism & Phenotyping core in Baylor College of Medicine at Houston, Texas. Insulin, adiponectin and leptin were measured by ELISA (Millipore) following manufacturer protocol.

### Luminex detection of proinflammatory cytokines

Serum cytokines were detected with magnetic bead technology (Invitrogen, Cytokine & Chemokine 26-Plex Mouse ProcartaPlex™ PanelLuminex) following manufacturer instructions.

### RNA extraction and quantitative RT-PCR analysis

#### RNA extraction and reverse transcription

Cultured T cells were homogenised in TRIzol® reagent (Thermo Fisher Scientific). An aqueous (RNA-containing) phase was generated using 1:5 bromo-chloro-propane, mixed 1:2 with 70% isopropanol and centrifuged at 12,000 g to precipitate RNA. Samples were treated with DNaseI (Qiagen). RNA concentration and integrity were determined by a NanoDrop™ One spectrophotometer (Thermo Scientific). Total RNA with an A260/A280 ratio ranging from 1.8 to 2.2 was converted to cDNA using the Maxima First Strand cDNA Synthesis Kit for RT-qPCR with dsDNase (Thermo Fisher Scientific).

#### Quantitative PCR

Quantitative PCR (qPCR) primers (Invitrogen-ThermoFisher) were designed using Primer-BLAST (NCBI; sequences are detailed below). A total of 10 ng of cDNA was used for quantitative PCR in a total volume of 10 µl with GoTaq® qPCR Master Mix (Promega) and specific primers, on a Bio-Rad CFX Opus 384 (Bio-Rad Laboratories). The amplification conditions were determined by the primers to present amplification efficiency close to 100% and a single peak in melt-curve analyses. Each Real-time PCR reaction was performed in triplicate. Glyceraldehyde 3-phospate dehydrogenase (*Gapdh* mRNA) and β-actin (*Actb* mRNA), encoding housekeeping proteins GAPDH and β-Actin, respectively, were included to monitor differences in RNA abundance. The log-fold change in mRNA expression was calculated from ΔΔCt values relative to control samples (T cells with acute stimulation and maintained with IL-2).

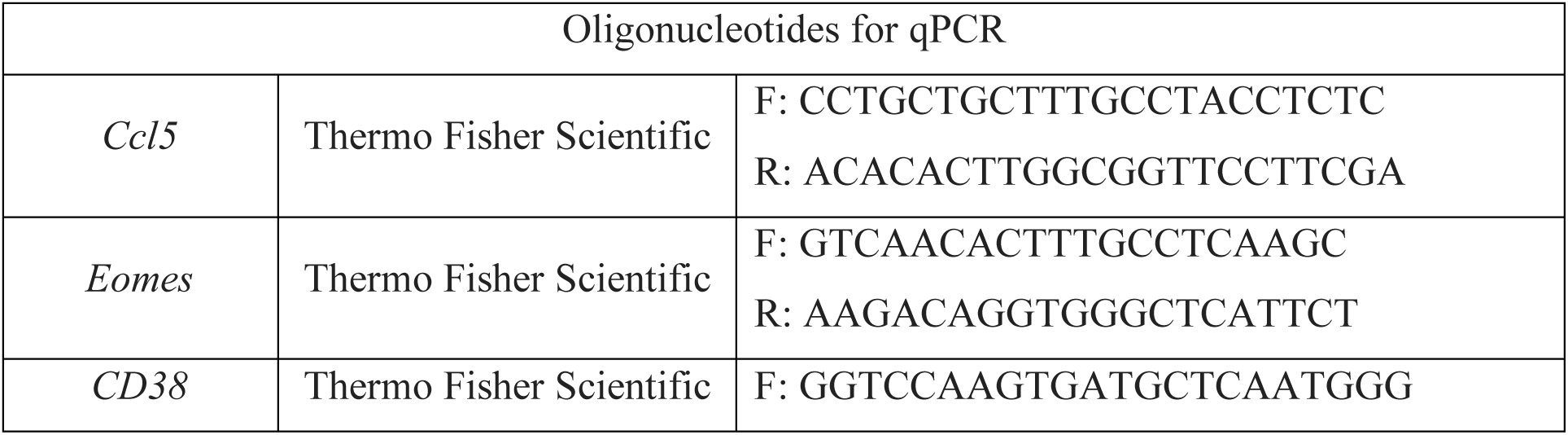
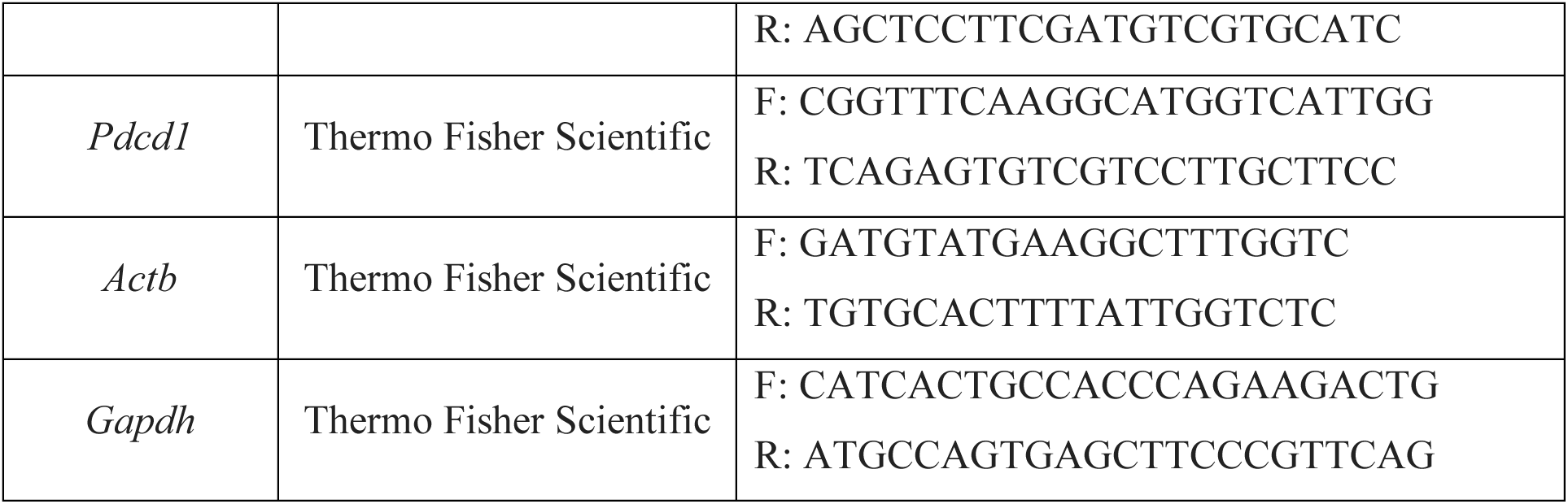

### Statistical analysis

Statistical analysis was performed with GraphPad Prism (version 9). Data are expressed as mean ± s.e.m. (standard error of the mean). Outliers were excluded by the ROUT method (5%). Comparisons for two groups were calculated using unpaired two-tailed Student’s *t*-tests (for two groups meeting the normal distribution criteria) or Mann-Whitney U test (for two groups without normal distribution) according to the Shapiro– Wilk normality test. Comparisons for more than three groups were calculated using One-Way ANOVA (for three or more groups meeting the normal distribution criteria) or Kruskal-Wallis test (for three or more groups without normal distribution) according to the Shapiro–Wilk normality test.

Unless otherwise specified, *n* represents the number of individual biological replicates and is represented in graphs as one dot per sample. Flow cytometry plots are representative of at least three replicates. No statistical method was used to predetermine sample size, but a minimum of three samples were used per experimental group and condition.

## Legends to supplementary material

**Extended Data Table 1. Statistic comparison of Extended Data Figure 6b,d**

**Extended Data Table 2. Statistic comparison of Figure 2g**

**Extended Data Figure 1.**
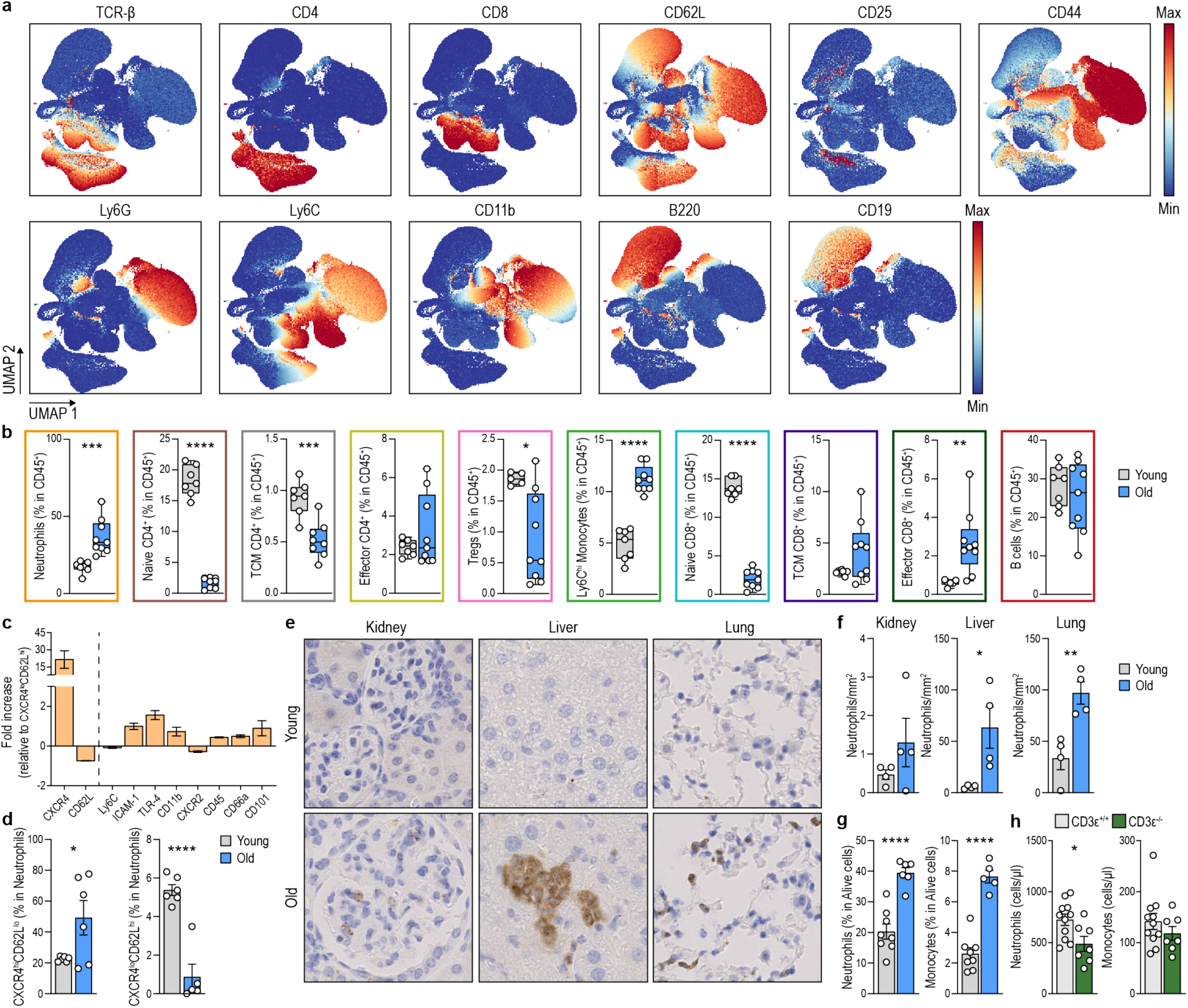
Enhanced granulopoiesis and accumulation of CXCR4^hi^CD62L^lo^ neutrophils in tissues from aged mice. **a**, UMAP representation of the expression levels of markers used to identify the different populations of blood cells shown in Fig. 1a and Extended Fig. 7c. **b**, Boxplots comparing the percentage of the different clusters in blood from young (3 mo) and old (24 mo) mice (n = 6-9 mice per group). **c**, Relative CXCR4^lo^ CD62L^hi^ to CXCR4^hi^ CD62L^lo^ expression of different surface markers showing an upregulation of pro-inflammatory markers in CXCR4^hi^ CD62L^lo^ neutrophils (n = 6-16 mice per group). **d**, Quantification of the percentage of CXCR4^hi^ CD62L^lo^ and CXCR4^lo^ CD62L^hi^ neutrophils in spleen from young (3 months old, mo) and old (24 mo) mice (n = 6 mice per group). **e,f**, Representative immunohistochemistry images (**e**) and quantification (**f**) of myeloperoxidase-positive neutrophils in kidney, liver, and lung sections from young (3 mo) and old (24 mo) mice (n = 4 mice per group). **g**, Percentage of neutrophils and Ly6C^hi^ monocytes in the BM from young (3 mo) and old (24 mo) mice assessed by flow cytometry (n = 6-8 mice per group). **h**, Percentage of neutrophils and Ly6C^hi^ monocytes in blood from old (17 mo) T cell deficient *Cd3e*^-/-^ and control mice (n = 7-11 mice per group).

**Extended Data Figure 2.**
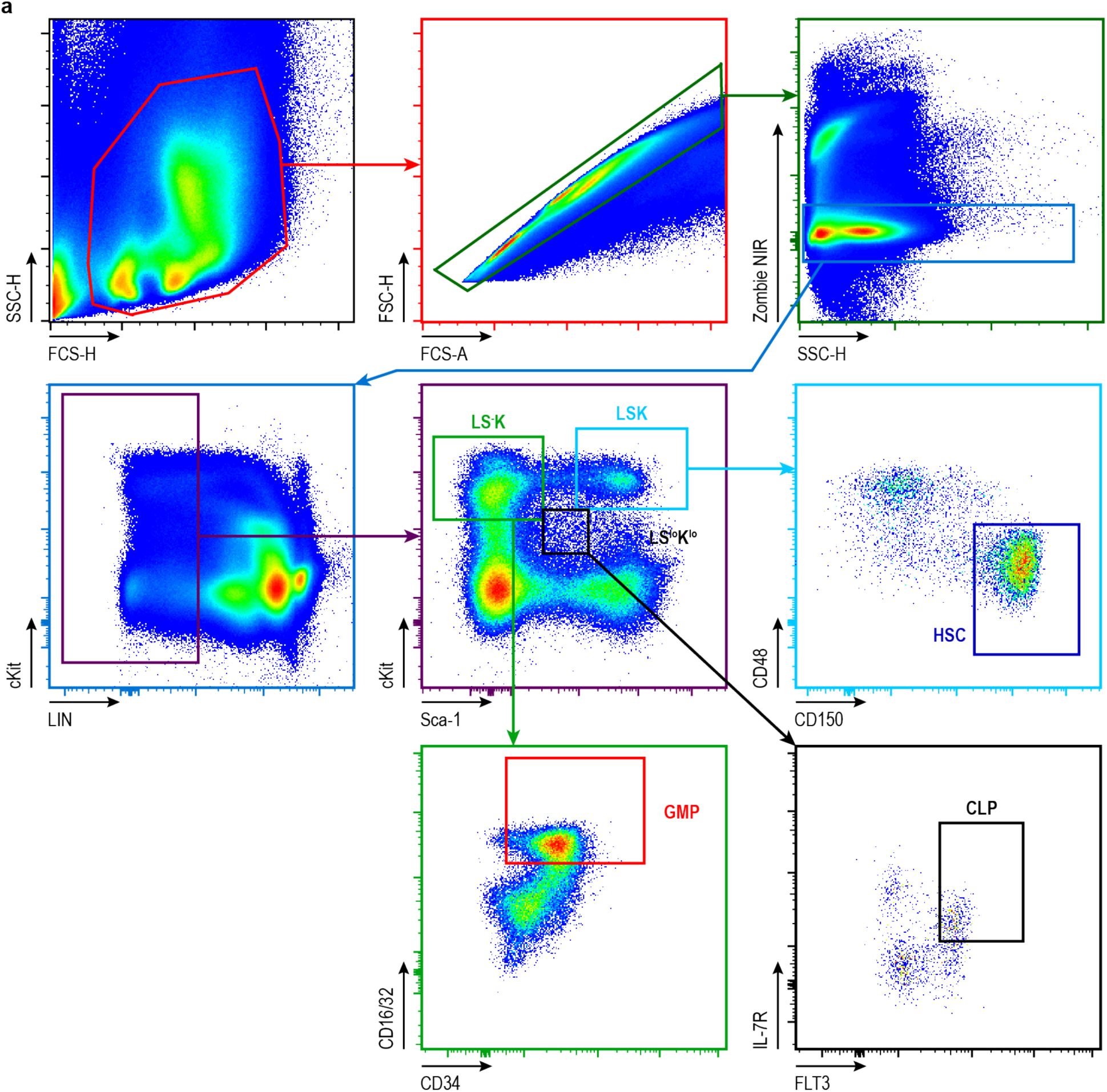
Gating strategy to identify hematopoietic precursors in the bone marrow. **a**, Gating strategy followed to identify LS^-^K (Lineage^-^Sca^-^cKit^+^) cells, GMPs (LS^-^K; CD16/32^+^CD34^+^), LSK (Lineage^-^Sca^+^cKit^+^) cells, HSCs (LSK; CD48^+^CD150^-^), LS^lo^K^lo^ (Lineage^-^Sca^lo^cKit^lo^) cells and CLPs (LS^lo^K^lo^; IL-7R^+^FLT3^+^).

**Extended Data Figure 3.**
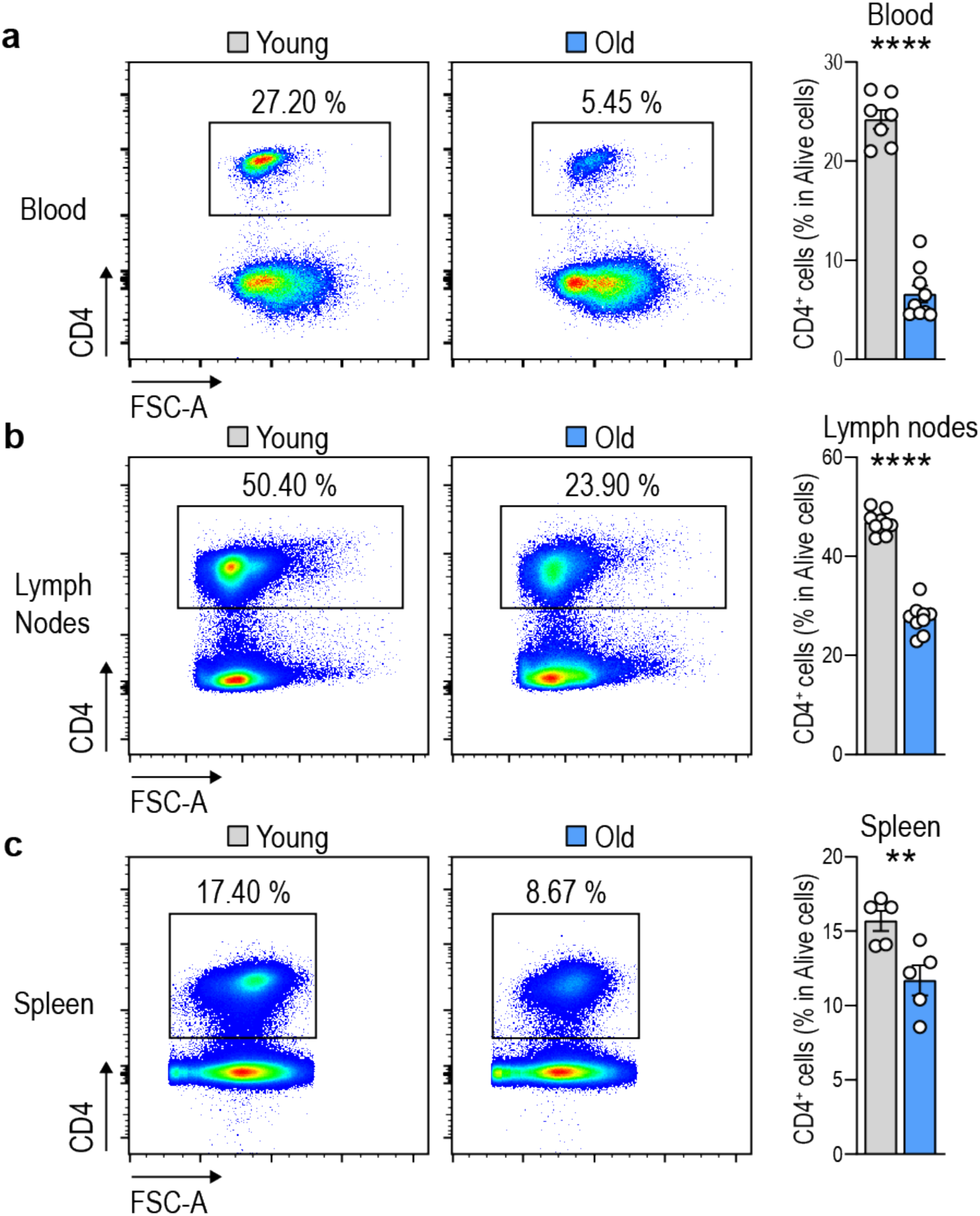
Decreased percentage of CD4^+^ T cells in the blood, lymph nodes and spleen during ageing. **a-c**, Representative flow cytometry plots (left) and quantification (right) of the percentage of CD4^+^ cells in the blood (**a**), lymph nodes (**b**) and spleen (**c**) from young (3 mo) and old (24 mo) mice (n = 5-9 mice per group).

**Extended Data Figure 4.**
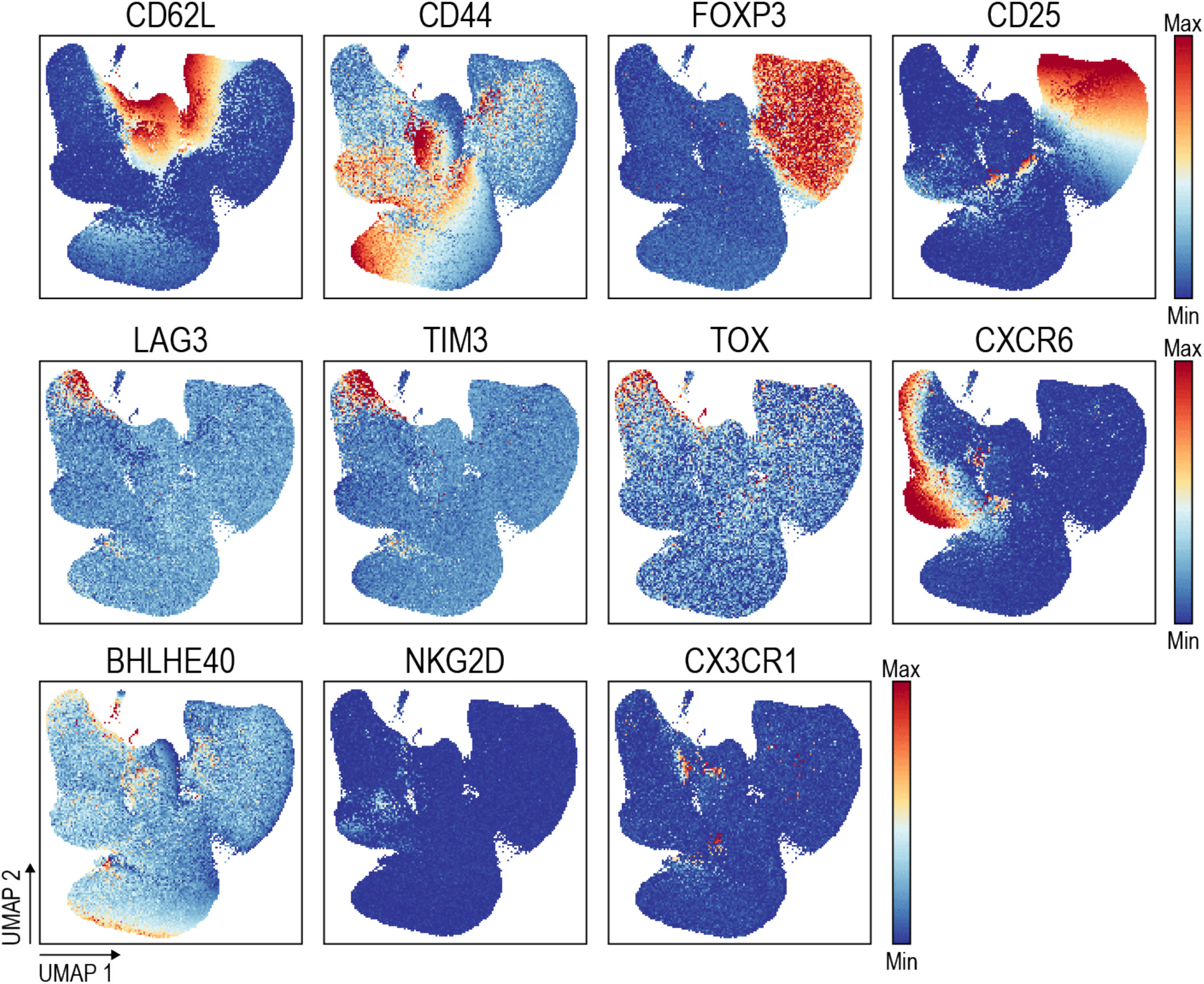
Expression of markers identifying CD4^+^ clusters in the BM. UMAP representation of the expression levels of markers used to identify the different populations of BM CD4^+^ cells shown in Fig. 2a.

**Extended Data Figure 5.**
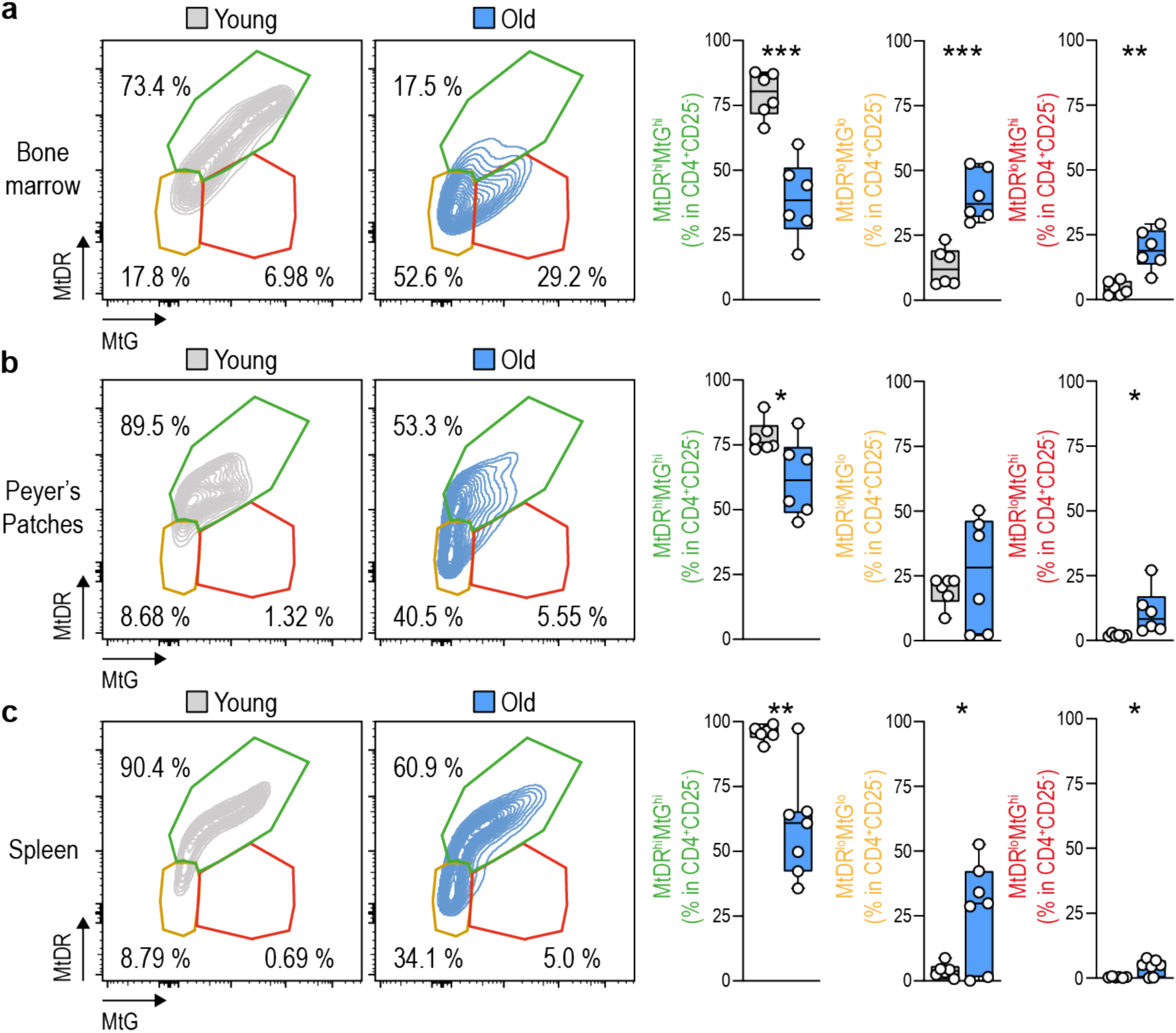
CD4^+^ T cells with dysfunctional mitochondria preferentially accumulate in the BM. **a**-**c,** Representative contour plots (right) and quantification of the percentage of CD4^+^ cells exhibiting polarized (MtDR^hi^MtG^lo^) and depolarized (MtDR^lo^MtG^lo^ or MtDR^lo^MtG^hi^) mitochondria in the BM (**a**), Peyer’s Patches (**b**) and the spleen (**c**) from young (3 mo) and old (24 mo) mice (n = 6 mice per group).

**Extended Data Figure 6.**
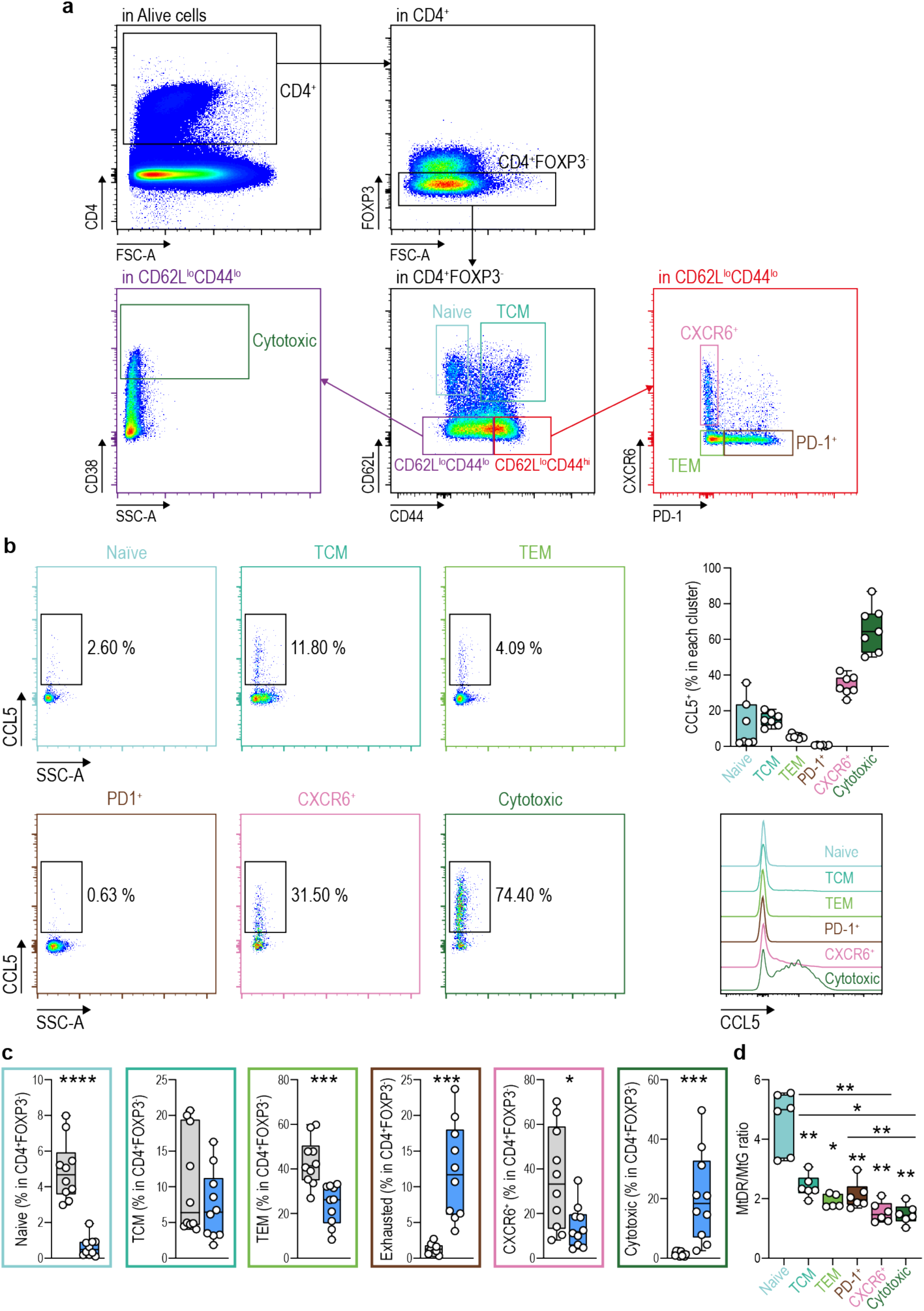
Identification of CD4^+^ CTLs by surface markers. **a**, Intracellular staining demonstrating the successful identification of all T cell clusters identified in the BM by extracellular staining. T regulatory cells were excluded by gating on CD4^+^Foxp3^-^ cells (for extracellular staining Foxp3 was substituted by CD25). First, Naïve (CD4^+^CD62L^hi^CD44^-^) and TCM (CD4^+^CD62L^hi^CD44^+^) cells were identified. Then, CD44^hi^CD62L^-^ cells or CD44^lo^CD62L^-^ were gated. CXCR6^+^ (CD4^+^CD62L^-^ CD44^hi^PD-1^-^CXCR6^hi)^, PD-1^+^ (CD4^+^CD62L^-^CD44^hi^CXCR6^-^PD-1^+^) and TEM (CD4^+^CD62L^-^CD44^hi^CXCR6^-^PD-1^-^) cells were identified in the CD44^hi^CD62L^-^ population. CD4^+^ CTLs were identified as CD4^+^CD62L^-^CD44^lo^CD38^hi^. **b**, Representative flow cytometry staining for CCL5 in all identified clusters. A high enrichment of CCL5^+^ cells is observed in the population corresponding to CD4^+^ CTLs. (**right up**), Boxplots showing the quantification of the percentage of CD4^+^ CCL5^+^ cells included on each cluster when identified by extracellular multiparametric spectral flow cytometry staining in BM samples from old (24 months old, mo) mice (n = 10 mice per group). (**right down**), Representative histogram showing the expression of CCL5 in the different clusters (n = 7 mice per group). **c**, Boxplots showing the quantification of the percentage of each cluster in CD4^+^Foxp3^-^ cells identified by extracellular multiparametric spectral flow cytometry staining in BM samples from young (3 mo) and old (24 mo) mice (n = 10 mice per group). The frequencies and percentages of cells on each cluster are similar to the observed by intracellular staining. **d**, MtDR/MtG geometric mean fluorescence intensity ratio in all identified clusters in BM samples from old mice (n = 6 mice per group).

**Extended Data Figure 7.**
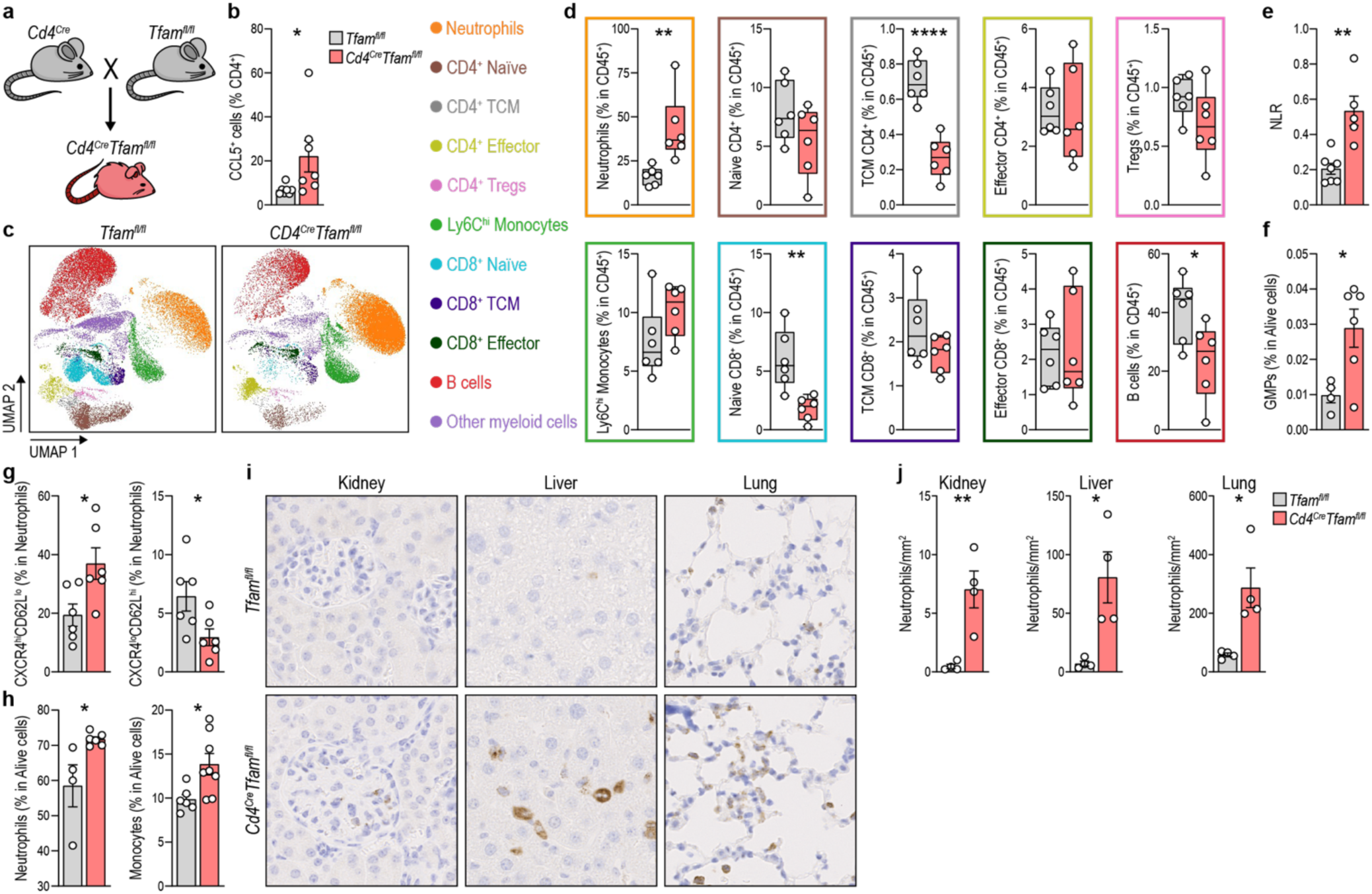
Genetic induction of a mitochondrial dysfunction in T cells enhances granulopoiesis and neutrophil output in mice. **a**, Diagram showing the genetic strategy followed to generate *CD4^Cre^ Tfam^fl/fl^* mice. **b**, Percentage of splenic CD4^+^CCL5^+^ cells in *Tfam^fl/fl^*and *CD4^Cre^ Tfam^fl/fl^* mice (n = 7 mice per group). **c**, UMAP with Cluster-X overlay showing the distribution of the eleven clusters of immune cells identified by multiparametric spectral flow cytometry in the circulation of *Tfam^fl/fl^* (8 months old, mo) and *CD4^Cre^ Tfam^fl/fl^* (8 mo) mice. **d**, Boxplots comparing the percentage of the different depicted populations in *Tfam^fl/fl^* and *CD4^Cre^ Tfam^fl/fl^* mice (n = 6 mice per group). **e**, Quantification of the NLR in the circulation of *Tfam^fl/fl^* and *CD4^Cre^ Tfam^fl/fl^* mice (n = 5-7 mice per group). **f**, Percentage of GMPs in the BM of *Tfam^fl/fl^*and *CD4^Cre^ Tfam^fl/fl^* mice (n = 4-6 mice per group). **g**, Quantification of the percentage of CXCR4^hi^CD62L^lo^ and CXCR4^lo^CD62L^hi^ neutrophils in blood from of *Tfam^fl/fl^* and *CD4^Cre^ Tfam^fl/fl^* mice (n = 4-6 mice per group). **h**, Percentage of neutrophils and Ly6C^hi^ monocytes in the BM of *Tfam^fl/fl^*and *CD4^Cre^ Tfam^fl/fl^* mice (n = 4-6 mice per group). **i**, Immunohistochemistry for MPO in kidney, liver and lung sections from *Tfam^fl/fl^*and *CD4^Cre^ Tfam^fl/fl^* mice (n = 4 mice per group). **j**, Quantification of myeloperoxidase-positive neutrophils per area.

**Extended Data Figure 8.**
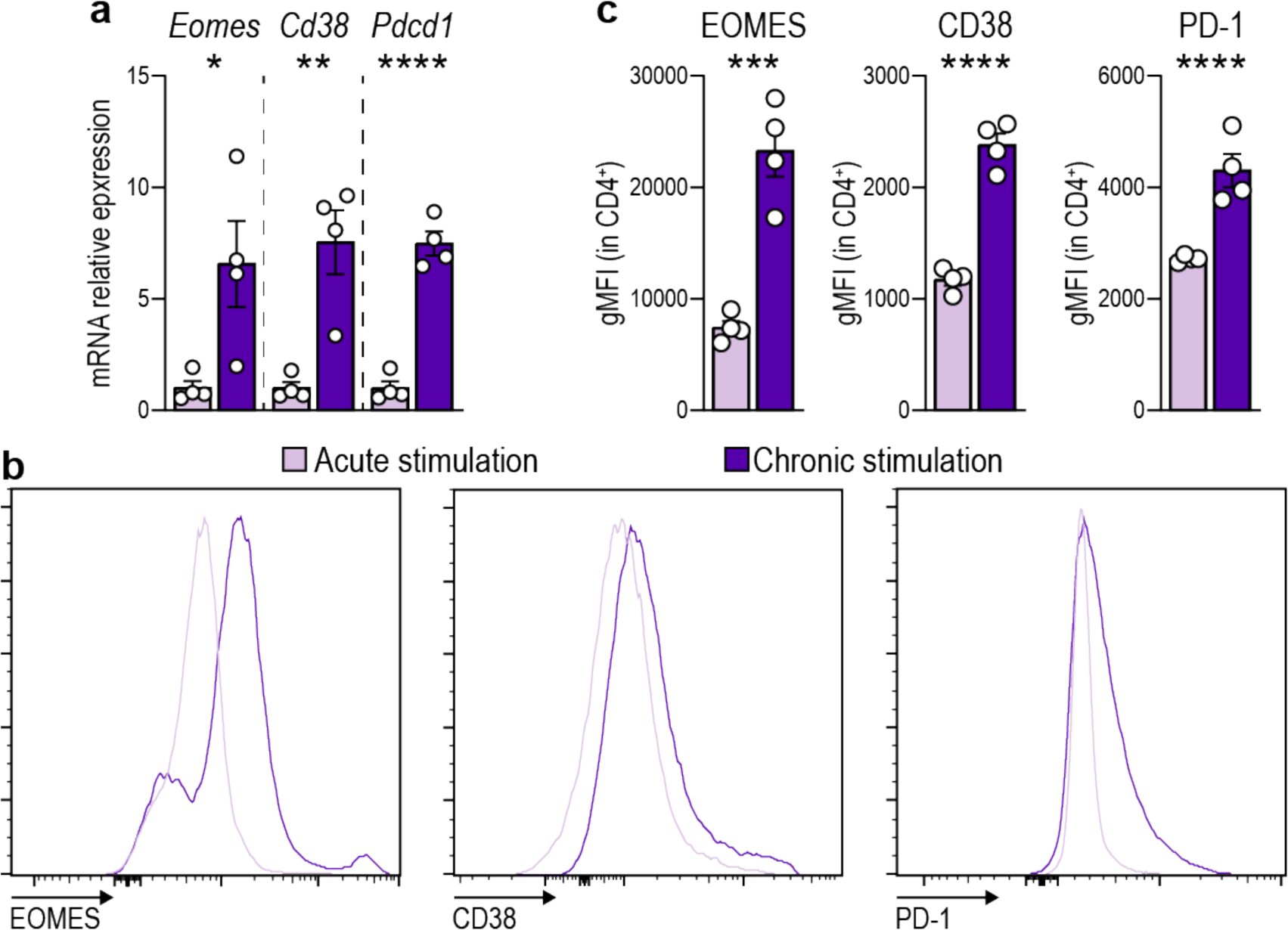
Prolonged TCR stimulation induces the expression of CD4^+^CTL markers in vitro. **a-c**, mRNA (**a**) and protein (**b,c**) levels of EOMES, CD38 and PD-1 in T cells acutely or chronically stimulated *in vitro* with anti-CD3/anti-CD28 determined by RT-qPCR analysis and flow cytometry, respectively (n = 4).

